# Connectivity underlying motor cortex activity during naturalistic goal-directed behavior

**DOI:** 10.1101/2023.11.25.568673

**Authors:** Arseny Finkelstein, Kayvon Daie, Márton Rózsa, Ran Darshan, Karel Svoboda

## Abstract

Neural representations of information are shaped by local network interactions. Previous studies linking neural coding and cortical connectivity focused on stimulus selectivity in the sensory cortex ^1–4^. Here we study neural activity in the motor cortex during naturalistic behavior in which mice gathered rewards with multidirectional tongue reaching. This behavior does not require training and thus allowed us to probe neural coding and connectivity in motor cortex before its activity is shaped by learning a specific task. Neurons typically responded during and after reaching movements and exhibited conjunctive tuning to target location and reward outcome. We used an all-optical ^5,4,6,7^ method for large-scale causal functional connectivity mapping *in vivo*. Mapping connectivity between > 20,000,000 excitatory neuronal pairs revealed fine-scale columnar architecture in layer 2/3 of the motor cortex. Neurons displayed local (< 100 µm) like-to-like connectivity according to target-location tuning, and inhibition over longer spatial scales. Connectivity patterns comprised a continuum, with abundant weakly connected neurons and sparse strongly connected neurons that function as network hubs. Hub neurons were weakly tuned to target-location and reward-outcome but strongly influenced neighboring neurons. This network of neurons, encoding location and outcome of movements to different motor goals, may be a general substrate for rapid learning of complex, goal-directed behaviors.

## Introduction

A fundamental challenge in brain research is understanding how connections between neurons shape the activity patterns underlying neural computation and behavior. During behavior, intermingled cortical neurons exhibit complex and diverse dynamics ^8–14,6,15^. Each neuron receives inputs from hundreds of local neurons ^16,17^ and long-range inputs ^18–20^, and the strengths of these inputs can be modulated *in vivo* ^21,22,14,23,24^. Therefore, understanding the relationship between connectivity and activity requires measuring behavior-related dynamics of individual neurons and mapping connectivity between the same neurons *in vivo*.

The motor cortex plays a central role in orchestrating goal-directed movements ^25^, and its neurons encode various movement parameters, including target location, muscle activation, effector velocity, and others. Motor cortex function has been studied in expert animals, trained to perform specific behavioral tasks, for example, skilled eye ^8^, arm ^26,10,9,12^, or tongue movements ^27,11,15^. However, motor cortex activity changes profoundly during learning to support skilled movements ^28,29,12,30^. We know relatively little about motor cortex activity before the network is shaped by task-specific learning over many trials.

To study neural coding in the motor cortex before learning reshaped neural dynamics and the underlying connectivity, we developed a naturalistic behavior that does not require training, in which mice gathered reward in one of many locations with multidirectional tongue reaching. We focused on the anterior lateral motor cortex (ALM) – an area critical for the planning and execution of sensory-guided licking in mice ^11,31,13,6,14,32,33^. Cellular calcium imaging revealed ALM neurons with exquisite tuning to target-location and reward-outcome. We used all-optical methods to measure the influence of individual neurons on activity in other neurons (‘causal connectivity’) ^5,4,6,7^. These causal connectivity maps uncovered sparse, highly connected hub neurons embedded within a fine-scale columnar architecture with like-to-like connectivity between target-location neurons.

## Results

### Neural correlates of multidirectional tongue-reaching

Head-fixed mice were presented with a port (target) that they could lick for a water reward. On each trial, the target was presented at one out of multiple (9, 16, or 25) possible locations in front of the mouse face (**Fig. 1a**). The target was moved outside the mouse’s reach between trials. Mice quickly became engaged in the task and reached for water rewards with their tongue (**Fig. 1b, c**), typically within the first behavioral session, and performed hundreds of trials per session (**Extended Data Fig. 1**).

**Fig. 1.**
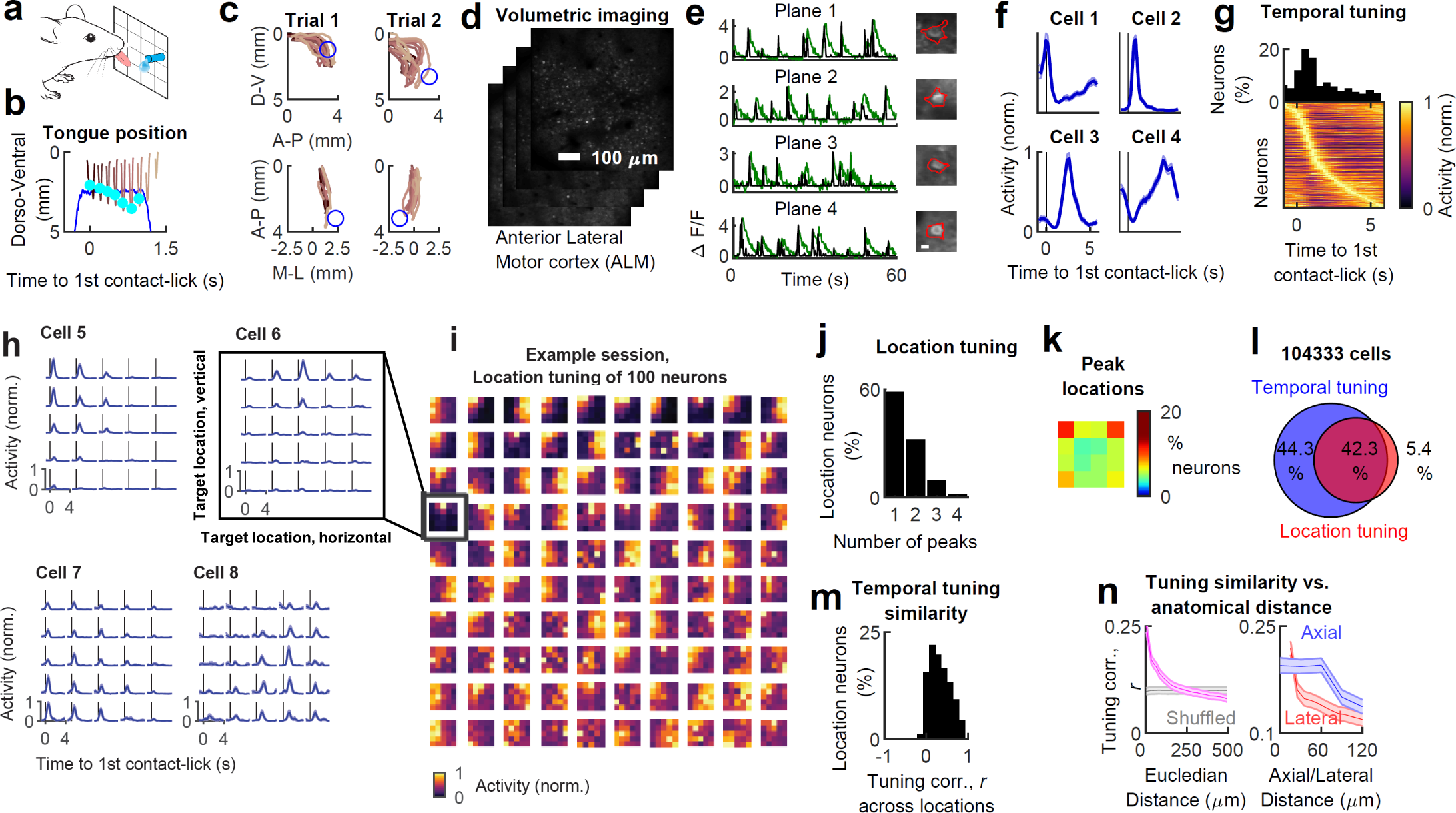
Neural activity in ALM during tongue-reaching. **a,** Schematic showing a grid of possible target locations. The mouse licks the target (spout) for a water reward. **b**, The trial begins with the target ascending to a reachable location. Tongue (brown) and target (blue) trajectories are shown for a single trial; cyan circles indicate tongue contact with the target. **c**, 2-D views of tongue trajectories (brown) towards two different target locations (blue) in two trials. Doral-ventral (D-V), anterior-posterior (A-P), and medial-lateral (M-L) projections are shown. **d**, Volumetric calcium imaging showing four planes imaged in ALM layer 2/3 across 120 µm depth. **e**, ΔF/F traces (green) and deconvolved spike trains (black) from four example cells (right, red delineates individual cells) imaged in different planes; scalebar, 10µm. **f**, Temporal tuning of four example cells, averaged across trials. **g**, Bottom, neural activity of all neurons with significant temporal tuning; each row corresponds to the trial-averaged response of one neuron (of 90,336). Top, preferred response time of all neurons. **h**, Location-temporal tuning of four example cells; traces show trial-averaged responses in each target location. **i**, Target location tuning of 100 simultaneously imaged cells. **j-k**, Distribution of number of peaks (j) and peak locations (k) in location maps of location-tuned neurons. **l,** Proportions of neurons with temporal and locational tuning out of all imaged cells in ALM. **m**, Similarity in temporal tuning across different target locations, for location-tuned neurons. **n**, Tuning similarity (of location-temporal tuning; Methods) between pairs of neurons as a function of their anatomical distance. Data in f-h is displayed as mean ± s.e.m. across trials. Data in n is displayed as mean ± s.e.m. across sessions.

We measured neural activity using volumetric calcium imaging of excitatory neurons in layer 2/3 of ALM (**Fig. 1d-e**) in mice expressing the calcium indicator GCaMP6s ^6,34^ (Methods). In all analyses, we refer to the deconvolved spike rate ^35,36^ as ‘activity’ (**Fig. 1e**; ΔF/F, green; deconvolved spike trains, black). Out of 104,333 imaged neurons, 86.6% (90,336 neurons) showed significant task-related activity modulation at different times during the trials (**Fig. 1f**; ‘*temporal tuning*’; **Extended Data Fig. 2a**). Similar to previous measurements in the motor cortex of untrained mice ^29,12^ and in contrast to trained animals ^8,26,29,12,11,31,33^, neural activity in most neurons peaked after movement onset, sometimes seconds later (**Fig. 1g**).

Neurons carried information about target location (47.7 %), with elevated activity for particular target locations (**Fig. 1h-i**, ‘*location tuning*’). Most neurons had unimodal tuning – with a single target location associated with a peak of activity (**Fig. 1j**; ‘number of peaks’; Methods), with peaks ranging from broad to narrow (**Extended Data Fig. 2b-c**). Across neurons, peaks of activity spanned the entire range of target locations but were biased towards the upper corners of the grid (**Fig. 1k)**. Notably, ALM neurons displayed tuning from the first day of behavior (**Extended Data Fig. 2d**), indicating that target location tuning reflects neural coding for naturalistic goal-directed reaching.

Location-tuned neurons were active at specific times during the behavior (**Fig. 1h,l**; 88.6% of location tuned neurons had temporal tuning), and temporal and location tuning were reliable across trials (**Extended Data Fig. 2e-h**). Location-tuned neurons had similar temporal tuning across all target locations (**Fig. 1m**), suggesting conjunctive encoding of location and response time. The peak-response times of different location-tuned neurons spanned the entire duration of the trial (**Extended Data Fig. 2i**), but this distribution was not uniform. Specifically, 37.3% of location-tuned neurons responded within one second after the first tongue-target contact, consistent with movement-related activity. Interestingly, almost ∼30% of these neurons responded much later (2-6 s after first contact; typical licking duration was <2 s, **Extended Data Fig. 1b**), thus carrying information about the target location after the movement ended.

We probed the spatial organization of neurons as a function of their combined temporal and location tuning (Methods; the results were similar when location and temporal tunings were analyzed independently **Extended Data Fig. 2j,k**). Neurons with different tuning appeared intermingled on the scale of > 250 µm (**Fig. 1n**, left). However, similarly tuned neurons were likelier to be near each other on smaller spatial scales (**Fig. 1n**, left). This similarity in tuning decayed faster laterally than axially (dorsal-ventral) (**Fig. 1n**, right), indicating that neurons with similar tuning were organized in columns.

Internal monitoring of action outcomes is fundamental for goal-directed behavior and learning ^37–42^. We next asked whether ALM neurons carry information about the outcome of tongue-reaching to different target locations. We randomly increased or omitted rewards on 20% of trials (**Fig. 2a**). Many neurons were significantly modulated by reward increase (19.5%) or omission (10.0% of neurons). Some neurons increased their response following reward increase or omission (**Fig. 2b**, cell 1-3,6), whereas others suppressed their response (**Fig. 2b**, cell 4-5). This ‘*reward-outcome tuning*’ was not explained by differences in licking for different reward sizes (**Extended Data Fig. 3**). Overall, 13.8 % of ALM neurons showed conjunctive tuning to target location and reward outcome (**Fig. 2c-d**).

**Fig. 2.**
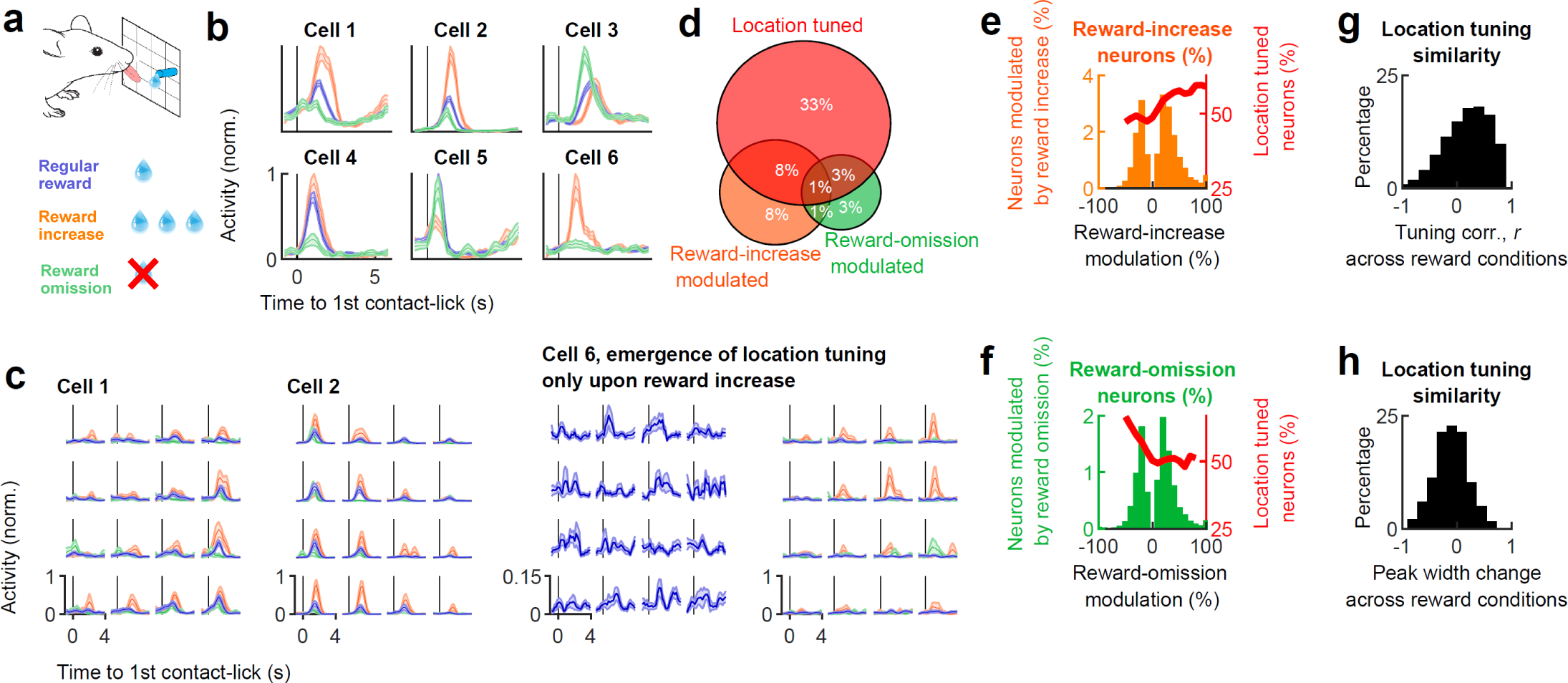
Conjunctive tuning to target location and reward-outcome. **a**, Reward was randomly increased or omitted on 20% of the trials. **b**, Temporal tuning of six examples cells that are modulated by reward outcome, averaged across trials with regular reward (blue), reward increase (orange), or reward omission (green). **c**, Target location tuning of three example cells with reward outcome modulation. Cells 1, 2, 6 are the same cells shown in b. **d,** Proportions of neurons with reward-outcome and/or location tuning in ALM. **e-f**, Percentage of neurons (left y-axis) with different levels of reward modulation, in neurons with modulation by reward-increase (e) or reward omission (f). Percentage of location-tuned neurons (right y-axis) among neurons with different levels of reward-modulation. **g-h** Similarities in location tuning shape (g) and peak width (h) across different reward conditions, in neurons with conjunctive tuning to location and reward outcome. Data in b-c is displayed as mean ± s.e.m. across trials.

The fraction of location-tuned neurons was higher among neurons up-modulated by reward increase (**Fig. 2e)** or among neurons suppressed by reward omission (**Fig. 2f)**, implying that the activity of location-tuned neurons tends to increase with more rewarding (positive) outcomes. In most neurons with conjunctive tuning (location × reward-outcome), the reward outcome modulated the amplitude but not the shape of the location tuning (measured by the correlations in location tuning and the peak width; **Fig. 2g-h**), suggesting that reward-related information acted as a gain modulation of location tuning. Notably, in a few cells (1% < of total cells), which did not display location tuning under regular reward conditions, sharp location tuning ‘emerged’ upon reward increase (**Fig. 2c**, cell 6). Thus, as a population, ALM neurons encoded the consequence of movements towards different target locations.

### Optical mapping of causal connectivity

Anatomical clustering of neurons with similar tuning (**Fig. 1n**) can arise from local connectivity ^27,2^. To test the relationship between network connectivity and the tuning of individual ALM neurons during behavior, we developed a method for rapid measurement of causal connectivity between excitatory neurons ^4^. Imaged neurons were randomly selected for two-photon photostimulation (one target at a time) while evoked responses (suprathreshold; not necessarily monosynaptic) were measured in other excitatory neurons (**Fig. 3a**; Methods). For two-photon photostimulation, we used the light-activated soma-targeted (ST) cation channel ChrimsonR ^43^. Because motor cortex connectivity is directed from superficial layers to deeper layers ^44^, we photostimulated neurons in the most superficial imaging plane while randomly switching photostimulation targets every 192 ms (**Fig. 3b, Extended Data Fig. 4**; ∼ 150 target neurons per session). This randomization of photostimulation targets, combined with volumetric imaging, allowed rapid (30 mins) mapping of causal connections between ∼300,000 pairs of neurons within a 700×700×120 µm ALM volume (∼150 targeted × ∼2000 imaged neurons) per session. In each session, we imaged the activity of the same neurons during 1) baseline at rest, followed by 2) connectivity mapping at rest, and then 3) during multidirectional tongue-reaching behavior (**Fig. 3c**).

**Fig. 3.**
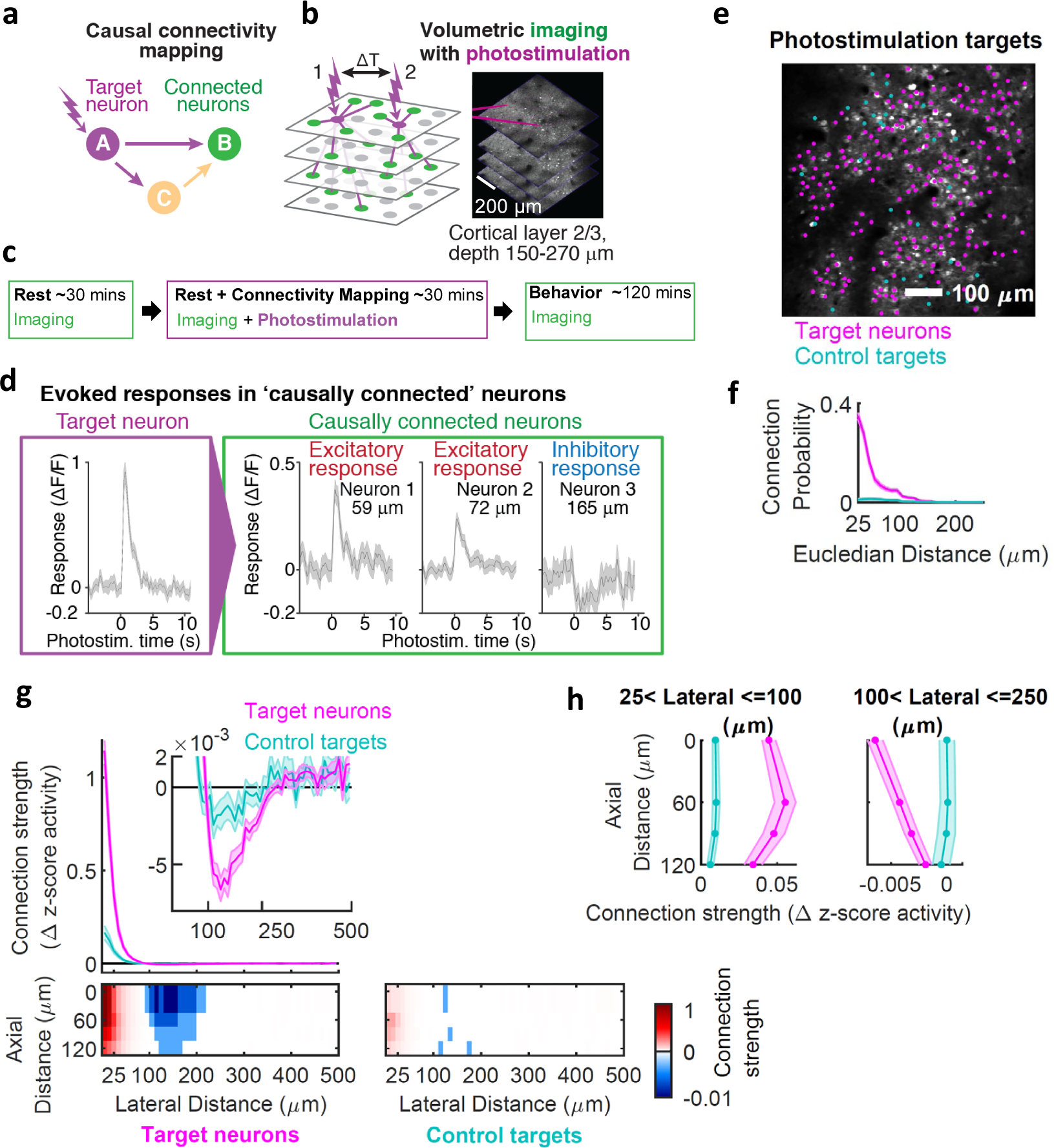
Mapping causal connectivity. **a,** Photostimulation of target neurons (magenta) and measurement of evoked responses in non-stimulated neurons (green). This method detects the causal effect of increase in activity of one neuron on another neuron. **b**, Excitatory cells were photostimulated in the superficial plane, while imaging responses of other excitatory cells within a volume of layer 2/3, motor cortex. **c,** Timeline for one experimental session. **d,** Example trial-averaged responses (∼50 trials) of a directly photostimulated neuron (‘target neuron’, left) and ‘causally connected neurons’ (not directly photostimulated, right) located at different lateral distances from the target neuron. This method can detect inhibitory (polysynaptic) causal connections between some of the neurons. **e,** Example field of view showing the top imaged plane with responding target neurons (magenta) and ‘control targets’ (neurons that were photostimulated but did not respond, cyan). **f,** Causal ‘connection probability’ as a function of distance (Euclidian) between all pairs of target neurons and causally connected neurons. **g-h,** Responses (causal ‘connection strength’; Methods) across all pairs as a function of anatomical distance from each target neuron. Marginal distribution of the connection strength as a function of lateral (g) and axial (h) distance from target neurons. Connection strength followed a Mexican-hat like profile of excitation and inhibition over anatomical space, with causal connectivity extending axially more than laterally. Data in f-h is shown for target neurons (magenta) and control targets (cyan), and displayed as mean ± s.e.m. across sessions.

A neuron that was ‘directly photostimulated’ (at lateral distance ≤ 15 µm from the photostimulation center) and responded to photostimulation is defined as ‘target neuron’ (**Fig. 3d**, Methods; ^6^). Neurons with photostimulation-evoked responses residing further away from the photostimulation center (at lateral distance > 25 µm) are defined as ‘causally connected’ ^6^ to the target neuron (**Fig. 3d**, 3 example causally connected neurons with significant excitatory or inhibitory responses). The amplitude of photostimulation-evoked responses in target and causally connected neurons correlated on single trials (**Extended Data Fig. 5a-d**). Furthermore, in trials in which photostimulation failed to trigger a response in target neurons, there was also no response in connected neurons (**Extended Data Fig. 5b-c**). In addition to causal excitatory connections, we detected inhibitory interactions between excitatory neurons (**Fig. 3d**, right), likely mediated by polysynaptic chains involving inhibitory interneurons ^45,46,4,47,48^. To assess the potential for photostimulation of neurons outside of the photostimulation plane (off-target), each connectivity-mapping session included ∼50 neurons that were photostimulated but did not respond. These ‘control targets’ (i.e., non-responding neurons) were intermingled with target neurons (i.e., with neurons that responded to photostimulation; **Extended Data Fig. 6a-b**). Thus, control targets still had neurons above or below, as well as neurites from nearby neurons in their vicinity, which could potentially respond to photostimulation.

We analyzed causal connectivity in networks of excitatory neurons (118,941 imaged neurons; 9,093 target neurons; putative connection pairs, > 22,000,000). The probability of an excitatory causal connection decayed quickly with distance from the target neuron (**Fig. 3f**, magenta), consistent with previous studies ^49,46,4^. Based on control targets, we estimated that off-target photostimulation contributed to approximately <10% of the responses in causally connected neurons (**Fig. 3f**, compare the magenta and cyan curves, corresponding to target neurons versus control targets; **Extended Data Fig. 6c**).

We next examined the three-dimensional spatial profile of excitation and inhibition evoked by photostimulation, averaged across all neuronal pairs. In the lateral dimension, excitation decayed within 100 µm (laterally) from the target neuron (**Fig. 3g** bottom, red). Further away (100-250 µm laterally), inhibition dominated (**Fig. 3g** bottom, blue). On average, these inhibitory interactions between excitatory neurons were two orders of magnitude weaker than excitatory interactions (see **Fig. 3g**, top; inset). That was likely because such inhibitory responses were polysynaptic (because we photostimulated and imaged excitatory neurons), and inhibition is harder to detect with GCaMP (because cytoplasmic calcium is already low for neurons with the low spike rates seen in layer 2/3 ^50^. In the axial dimension, excitatory responses peaked at 60 µm below the photostimulation plane (**Fig. 3h**, left) and extended for at least 120 µm (the deepest plane we imaged simultaneously with photostimulation). Excitation decayed faster laterally than axially (**Fig. 3h**, right). This effect could not be explained by off-target spread of photostimulation in the axial dimension because: i) The optical point spread function of the photostimulation was 11.7 µm axially ^6^, and ii) photostimulation of control targets showed much weaker responses (∼10 fold) compared to target neurons at all depths (**Fig.3g** bottom; **Fig. 3h**, left – compare magenta to cyan lines). This strong connectivity in the axial dimension could reflect preferential connections within ontogenetic radial clones of excitatory neurons ^51^. In contrast to excitation, inhibition peaked at the same plane as photostimulation and gradually decayed with depth (**Fig. 3h**, right). Taken together, causal connectivity in the motor cortex, similar to location tuning (**Fig. 1n)**, was also organized in functional columns, with a ‘Mexican hat’ profile of excitation and inhibition ^52,53^ over anatomical space.

### Relation between causal connectivity and functional organization

We next compared the functional properties of neurons and causal connectivity. We first analyzed the relationship between causal connectivity and location tuning. Neurons with stronger causal connectivity had more similar location tuning (**Fig. 4a**). This effect was apparent only for neurons residing within 100 µm from each other and could not be explained by distant-dependent modulation of tuning similarity alone (**Extended Data Fig. 7a**). We also found a relationship between connectivity strength measured during rest and ‘noise’ correlation measured either during rest or behavior, suggesting that causal connectivity mapping can explain network dynamics across different behavioral states (i.e., rest and behavior; **Extended Data Fig. 7b-e**). Taken together, neurons within a functional mini-column with similar location tuning were more likely to be connected.

**Fig. 4.**
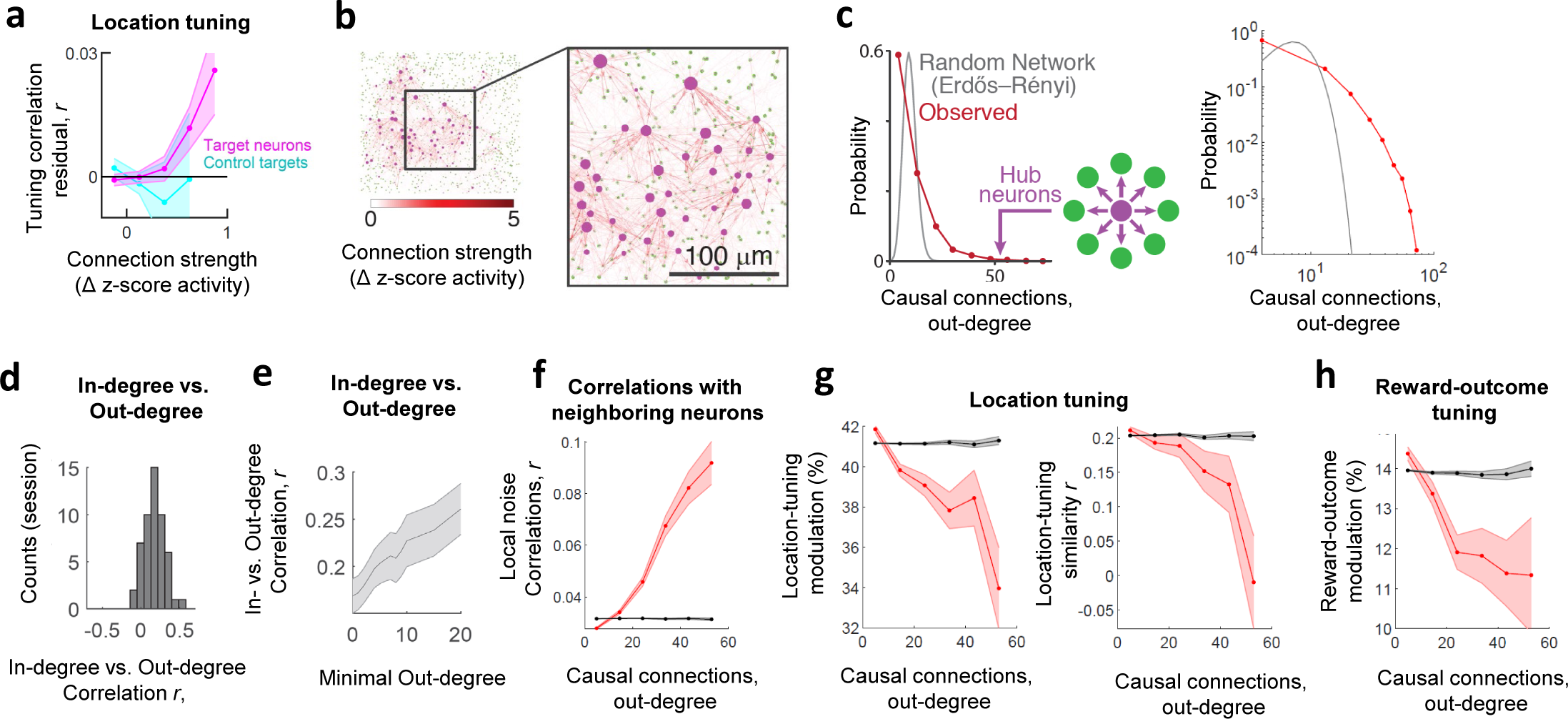
Causal connectivity in ALM. **a**, Comparison of causal connectivity strength with residual correlations in location tuning (Methods), shown for target neurons (magenta) and control targets (cyan) as mean ± s.e.m. across sessions. Neurons with similar location tuning were preferentially causally connected to each other**. b**, Example connectivity graph of the mouse motor cortex based on causal connectivity mapping across one session (top-view projection collapsed across all imaged planes). Node-size indicates the number of outgoing causal connections and arrow color the connection strength. Some neurons were much more connected than others. **c,** Distribution of the number of causal connections (out-degree) of targeted neurons shows a long tail (red), revealing hub neurons with a disproportionally high number of connections, inconsistent with random network connectivity (gray). **d,** Average correlation between in- and out-degree distribution of causal connections between target neurons, for different sessions. **e,** Correlation between in- and out-degree distribution of causal connections in different subnetworks of target neurons, restricted to neurons with a minimal out-degree connectivity (*m*), as a function of *m*, displayed as mean ± s.e.m. across sessions. **f-h,** The relation between number of causal outgoing connections (out-degree) of a neuron and its (f) noise correlation with neighboring neurons, or (g) its location-tuning or (h) reward-outcome tuning; displayed as mean ± s.e.m. across neuronal pairs in bins. Gray line corresponds to shuffle distribution. Data in c, f-h is based on all significant causal connection pairs (*n* = 65,390 pairs; Methods).

So far, we focused on the average pairwise relationship between causal connectivity and functional properties. However, connectivity maps were complex (**Fig. 4b**). We next analyzed the network-level organization of this local circuitry. Connectivity patterns between neurons were non-random: out-degree distribution of target neurons showed a long tail, including neurons with a very large number of out-degree connections, which we refer to as ‘hub neurons’ (**Fig. 4c**). There was a positive correlation between the number of in versus out degree connections (**Fig. 4d**; Methods). This correlation increased when restricting the analysis to subnetworks of neurons with a minimum out-degree connectivity (a parameter we increased progressively; **Fig. 4e**). In other words, subnetworks of neurons with higher out-degree connectivity also had higher in-degree connectivity and therefore were more interconnected – resembling a ‘rich club’ phenomenon predicted from *in silico* models of cortical connectivity ^54^.

Neurons with larger numbers of output connections are expected to have a greater influence on the local network. Indeed, we found that the activity of local neighboring neurons was more correlated to activity of neurons with more output connections (hubs; **Fig. 4f**). Finally, we asked whether neurons with different number of output connections differ in tuning properties. Hub neurons showed weaker location tuning (**Fig. 4g)** and weaker reward modulation (**Fig. 4h**) than less-connected neurons. Taken together, this suggests that i) local causal connectivity is not random, ii) connectivity predicts tuning, and iii) hub neurons are less tuned to task-related features but show more coupling to nearby neurons – suggesting that they may act as local coordinators of neural activity irrespective of the specific target location.

## Discussion

The motor cortex plays important roles in shaping goal-directed and skilled movements ^25^. Activity in descending pyramidal tract neurons helps initiate movements ^55,33^. However, neural dynamics in the motor cortex have been typically studied in mice trained on specific motor tasks. For instance, in mice trained in a memory-guided directional licking task, a large proportion of ALM neurons were selective for licking direction (left/right) before and during licking ^11,31,13–15^. More generally, in trained animals, modulation of motor cortex activity is most pronounced before and during movement ^26,8,9,29,12^.

In contrast to trained animals, we found that in naive mice, most ALM neurons showed task-related modulation after movement onset and even after movement. Our behavioral paradigm allowed to map neural responses to a continuum of target locations, similarly to studies of reaching and eye-movement in monkeys ^10,8,9^, albeit without the need for training. Neurons were tuned to target locations and modulated by reward size. In goal-directed behaviors, action selection is influenced by the expected outcome of specific actions. Neural activity in the parietal and the frontal cortex is correlated with action values ^38,40,39,56,57,41^. Updating the neural dynamics encoding action values may require *vectorial* error signals associated not just with reward values but also with *specific* actions ^42,58^. Conjunctive encoding of target location and reward outcome in ALM neurons (**Fig. 2**) signals deviations from expected value and the location in motor space where the action took place. This vectorial error signal may provide a teaching signal for learning complex goal-directed behaviors.

Local similarity in tuning could arise from similarity in long-range inputs and/or preferential local connectivity ^2,4,6^. Most studies relating connectivity to function focused on a few cells at a time. For example, *in vivo* calcium imaging has been combined with *post hoc* analysis of synaptic connectivity in brain slices ^59,2,60^ or via large-scale electron microscopy reconstructions of local circuits ^61,62^. Although these methods detect mono-synaptic connections, they have low throughput. Moreover, *ex vivo* methods probe connectivity in a quiescent network state, but the effect of one neuron on another depends on the activity of the network ^63,22,47,14,23^. Indeed, structural connectivity in *C. elegans* does not predict functional interactions ^7^, stressing the need to study connectivity *in vivo*. Instead, we developed an all-optical method based on two-photon optogenetics and calcium imaging ^5,4,6,7^ to map causal connectivity between neurons in the intact brain. This method is efficient in that it allows probing many pairwise connections in a single session, but cannot detect subthreshold connections and does not differentiate monosynaptic and multi-synaptic causal interactions.

Combining naturalistic behavior with causal connectivity mapping revealed a relationship between neuronal tuning to target locations and the organization of motor cortex local circuits. Tuning similarity of ALM neurons decayed with anatomical distance (**Fig. 1n**), and causal connectivity exhibited a Mexican hat profile of excitation and inhibition, also as a function of anatomical distance (**Fig. 3g**). In contrast to sensory cortex, where photostimulation mainly resulted in lateral inhibition ^45,4,48,47,64^, in the motor cortex, photostimulation elicited local excitation (**Fig. 3g**; ^6^), likely because the motor cortex operates closer to threshold (in a fluctuation-driven regime) compared to sensory cortex ^65^. Finally, ALM neurons with similar target location tuning were preferentially causally connected. The like-to-like causal connectivity motif between target location neurons was embedded within functional columns with a Mexican-hat profile of local excitation and longer-range inhibition. Such connectivity profiles are often used in continuous attractor network models to explain the neural mechanisms underlying neuronal tuning to continuous variables, such as orientation tuning in the visual cortex ^66^ and spatial tuning in the hippocampal formation ^67,53^. Learned discrete attractor dynamics drive both the selectivity and timing of dynamics in ALM neurons ^13,14^ and connected structures ^68,69^ in tasks with fixed discrete target location. Our work opens the venue for studying the network mechanisms (e.g., continuous attractors) underlying naturalistic behaviors involving a continuum of motor goals, before learning induces changes in connectivity and dynamics.

We identified heterogeneous ^70,49^ network interactions among excitatory neurons in layer 2/3. The distribution of out-degree connections was heavy-tailed, and included rare neurons with a large number of connections suggestive of network hubs. Hub neurons were identified in the structural connectome of *C. elegans* ^71,72^, in rodent hippocampus and neocortex during development ^73,74^, and predicted based on simulations of cortical columns ^54^. Future work will reveal how cortical neurons with different functional/network properties (e.g., hubs) map onto distinct cell types and what are the specific roles of different topological elements of motor cortex networks in driving network dynamics and behavior.

## Methods

### Mice

Data are from 15 mice of either sex (age at the beginning of experiments, 60–240 d). All mice were CamK2a-tTA (JAX, catalog no. 007004) × Ai94 (TITL-GCaMP6s) (JAX, catalog no. 024104) × slc17a7 IRES Cre (JAX, catalog no. 023527), providing widespread expression of GCaMP6s in excitatory cortical neurons. All procedures were in accordance with protocols approved by the Janelia Institutional Animal Care and Use Committee.

### Surgical procedures

Cranial window surgeries were performed as described previously ^75^. In brief, circular (diameter, 3 mm) craniotomies were centered over ALM (2.5 mm anterior and 1.5 mm lateral from Bregma). We expressed the soma-targeted opsin ST-ChrimsonR ^43^ in excitatory neurons by injecting a virus (10^12^ titer; AAV2/2 camKII-KV2.1-ChrimsonR-FusionRed; Addgene, plasmid #102771) into the craniotomy, 400 µm below the dura (5-10 sites, 20–30 nl each), centered within the craniotomy and spaced by ∼500 μm between injection sites. The craniotomy was covered by a cranial window composed of three layers of circular glass (total thickness 450 μm). The diameter of the bottom two layers was 2.5 mm. The top layer was 3 mm or 3.5 mm and rested on the skull. The window was cemented using cyanoacrylate glue and dental acrylic (Lang Dental). A customized headbar was attached using cyanoacrylate glue and dental cement.

### Behavior

After waiting 3-4 weeks to allow for viral gene expression, mice were placed on water restriction (1 ml/day) in a reverse light cycle room. Behavioral experiment commenced 3–7 days later. Before conducting the behavioral experiments, mice were head-fixed for the first time and placed under a microscope for a brief imaging session, in which the quality of the cranial window, the GCaMP6s signal, and the ability to detect photostimulation responses were assessed. These sessions, which did not involve a behavioral task, served as a habituation to the head-fixation procedure and the experimental conditions. Typically, the actual behavioral experiments started after a single habituation session.

During behavioral experiments, mice were head-fixed, and water rewards were delivered through a lickpot, referred to as the ‘target’. The target location was set using electric motors (Zaber). The entire behavior was controlled by a custom-programmed state machine (Bpod, Sanworks). To track orofacial movements, two CMOS cameras (Teledyne FLIR) viewed the face from the side and bottom under infrared (IR) light illumination (940 nm LED). High-speed videos were recorded at 250 Hz using BIAS software (IO rodeo, https://bitbucket.org/iorodeo/bias). To track tongue movements, we trained DeepLabCut^76^. Contacts with the target were detected using a custom-built lick detector ^11^.

The behavioral protocol involved mice licking a target that could appear at various locations around the mouse’s face along the medial-lateral (left-right) and dorsal-ventral (up-down) axes (**Fig. 1a**). Before the first behavioral session, which occurred either on the same day or the day following the habituation session, the mouse was introduced to the target for the first time. This was accomplished by initially presenting the mouse with a target directly in front of its mouth and releasing small drops of water. Mice began licking the target within 1-5 minutes of its first presentation. Once a mouse began licking, we mapped the maximal distance it could comfortably reach with its tongue by gradually moving the target away from the center along the medial-lateral and dorsal-ventral axes. We then divided the reachable area into a grid of possible target locations. We typically used a 4×4 grid of possible locations, but in some sessions, we used 3×3 or 5×5 grids. The grid spacing differed along the medial-lateral versus the dorsal-ventral axes, according to the maximum reachable distance along each axis. Mice could locate the target using their sense of smell and whisker touch to guide them to the target, to ensure the task was as naturalistic as possible, and to avoid the need for pretraining the mice on this specific task.

Each trial began with the target positioned out of reach (below the mouse face) for one second. It then ascended to a reachable location (at one of the locations on the target grid; movement time, ∼150 ms). The mouse was given time (up to ten seconds) to initiate contact with the target, and the target was withdrawn once the mouse collected the reward (one second following the first contact with the target). Rather than switching the target location every trial, the target appeared at the same location for several consecutive trials (‘block’, consisting typically of 7 trials of repeated locations) until switching to a new random location within the grid.

To increase the mouse motivation in contacting the target during the first session, the reward was dispensed at the beginning of the trial (‘auto-reward’) – and remained attached to the target until the mouse licked it. In the subsequent session, the ‘auto-reward’ feature was turned off, and the reward was dispensed only after the mouse contacted the target. In cases when mice lost motivation and stopped licking in the middle of the session, the ‘auto-reward’ feature was turned on. Upon change of target location (a new block), the first trial in a block was an auto-reward trial to assist the mouse in finding the new target location. After a few behavioral sessions, the ‘auto-reward’ feature was no longer necessary.

The ‘regular reward’ size was ∼2 µl of water (∼80% of the trials). In a random subset of trials, we either tripled the reward size (‘reward increase’, ∼10% of the trials) or omitted the reward (‘reward omission’, ∼10% of the trials). The first trial in a block was always a regular reward trial. We introduced reward increase and reward omission trials only from the second behavioral session after the mice had formed an estimate of the expected reward size. Individual behavioral sessions comprised 786 ± 39 trials (mean ± s.e.m; the longest session had ∼1429 trials) performed over ∼2 hours (**Extended Data Fig. 1**). In some sessions, we readjusted the maximal reachable distance (grid dimensions) at the beginning of the sessions. It could take up to 150 trials to readjust grid dimensions to ensure that the mouse licked for all target locations. Because of these readjustments, the first 150 trials were excluded from the analysis.

### Microscope

Two-photon imaging and photostimulation were performed using a customized microscope with a photostimulation path ^6^. The two-photon photostimulation path consisted of a 1040-nm pulsed laser (Fidelity 10, Coherent), a Pockels cell (Conoptics) for power modulation, and a pair of galvanometer mirrors (Cambridge, 6215H) for beam positioning. Imaging was done with a 920-nm pulsed laser (Chameleon Ultra II, Coherent) and a galvo-resonant scanner (Thorlabs). Volumetric imaging was performed using an electric tunable lens (Optotune) placed in the imaging path. Imaging and photostimulation were controlled by ScanImage (Vidrio). We applied an automatic online motion correction in both lateral and axial dimensions using ScanImage by taking a reference Z-stack of the imaged field of view at the beginning of each imaging session. The field of view was typically 738 × 688 μm^2^ (512 × 512 pixels^2^). Volumetric imaging was performed across four planes (typical relative depth of the planes was 0, 60, 90, 120 µm). Planes were imaged sequentially (33 ms per plane) by adjusting the electric tunable lens. We introduced a gap of 15 ms by blanking the acquisition between planes to allow the lens to settle, resulting in imaging rates of 5.2 Hz for the entire 4-plane volumes (**Extended Data Fig. 4**).

### Two-photon calcium imaging

Imaging and photostimulation were done in layer 2/3 of the anterolateral motor cortex (typically 150 µm from brain surface, range, 125–250 µm) ^6^. On each day (session), we imaged neurons from a different field of view in ALM (∼5 sessions per animal). Each session (*n* = 44 sessions) contained three epochs (**Fig. 3c**): a) Imaging of neural activity at rest before the start of behavior (‘rest’, ∼30 mins); b) imaging and photostimulation at rest (‘rest + connectivity mapping’, ∼30 mins); c) imaging during the multidirectional licking task (‘behavior’, ∼120 mins). In addition, there were 9 sessions with only ‘behavior’ epoch, and 8 sessions with only ‘rest + connectivity mapping’ epoch.

We used Suite2p ^36^ to identify neurons in the calcium imaging data, extract their fluorescence traces, and correct for neuropil contamination. Fluorescent traces of each neuron were baseline corrected (ΔF/F) by dividing the fluorescence trace of that neuron by the rolling max of the rolling min, using a 60 s time window. The fluorescent traces were deconvolved into spike trains based on the OASIS algorithm ^35^ in Suite2p. All analyses were done on the deconvolved spike rates. We refer to the deconvolved spike rate as ‘activity’ in all analyses (**Fig. 1e**; ΔF/F, green; deconvolved spike trains, black).

### Analysis of task-related neuronal tuning

To compute task-related neural activity, neural activity was parsed into behavioral trials and aligned to the first contact lick on each trial. For each neuron, *temporal tuning* (**Fig. 1f**) was defined as the trial-averaged neural activity using all ‘regular reward’ trials (see Methods section, ‘Behavior’). Activity was normalized to the peak of the trial-averaged neural activity in the time interval *t* = [-2,5] s, relative to the first contact lick. *Location tuning modulation* was defined as the normalized difference between the maximal and minimum of the trial-averaged neural activity in that time interval (**Extended Data Fig. 2a**):

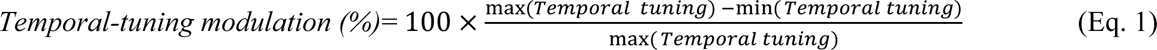

The time of the peak was defined as *peak response time*. We computed *temporal tuning stability* by splitting all trials (used to compute the tuning) into two independent sets of odd and even trials. We recomputed the tuning independently for odd and even trials and took the Pearson correlation coefficient (*r*) between the odd-trial and even-trial tuning curves as a measurement for tuning stability (**Extended Data Fig. 2f**). We defined the temporal tuning of a neuron to be significant for *r* (odd, even) ≥ 0.25 and location-tuning modulation ≥ 25%.

To compute temporal tuning in each target location (*location-temporal tuning*; **Fig. 1h**) for each neuron, we trial-averaged neural activity for all regular-reward trials in each target location. Neural activity was aligned to the first contact lick on each trial. Activity in each location bin was normalized to the *location-temporal peak* – the peak of trial-averaged neural activity in the bin with the highest activity in the time interval *t* = [-2, 5] s. We calculated location-temporal tuning stability by analogy to temporal tuning stability (see previous paragraph). We computed the temporal tuning in each location using two independent sets of trials (odd, even) and took the Pearson correlation coefficient (*r*) between the temporal tuning curves for each location bin, after concatenating across locations (**Extended Data Fig. 2g**). We defined the location-temporal tuning of a neuron to be significant for *r* (odd, even) ≥ 0.25.

To generate a *location-tuning map* for each neuron (**Fig. 1i**), we took the mean neural activity over the time interval *t* = [-2, 5] s. We averaged this value across trials in each location bin. The *peak location* of each neuron was the location bin corresponding to the maximum of the location tuning map. *Location tuning modulation* was defined as the normalized difference between the maximal and minimum of the location tuning map (**Extended Data Fig. 2b**):

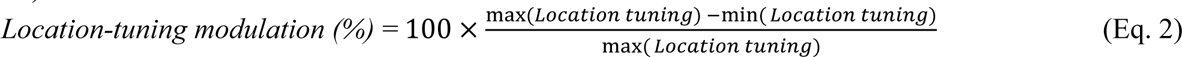

By analogy to temporal tuning stability (see previous paragraph), we calculated *location tuning stability*, by computing location tuning maps for two independent sets of trials (odd, even) and taking the Pearson correlation coefficient (*r*) between the two maps (**Extended Data Fig. 2h**). We defined the location tuning of a neuron to be significant for *r* (odd, even) ≥ 0.25 and location-tuning modulation ≥ 25%.

To compute the distribution of *peak locations* across the population (**Fig. 1k)**, we computed a two-dimensional histogram (expressed as a percentage of cells) of peak locations of all neurons with significant location tuning, imaged in experiments with a 4×4 location grid (24,654 neurons). A 4×4 location grid was chosen to ensure adequate location sampling, while having enough trials per location. The *number of peaks* (**Fig. 1j)** and *peak width* (**Extended Data Fig. 2c**) in the location tuning map of each neuron was also computed in experiments with a 4×4 location grid. To find the number of peaks, we scaled the location tuning map from 0 to 1 (i.e., from minimum to maximum activity). The *number of peaks* was defined as the number of local maxima with activity ≥ 0.5 (in the scaled map), which were separated from other local maxima by bins with activity ≤ 0.5. The *peak width* was defined also using the scaled map as the percentage of bins whose activity was ≥ 0.5, out of all bins.

To quantify all other metrics related to location tuning (**Fig. 1l-n**, **Fig. 2d-h, Extended Data Fig.2 b,g-i,k**), we combined experiments with different location-grid sizes (i.e., 3×3, 4×4, or 5×5) by resampling the behavioral trials according to a 3×3 grid. This allowed to 1) apply the same statistical criteria in all experiments, and 2) ensure that there are enough trials per location bin in all experimental conditions (e.g., across reward conditions). T*emporal tuning similarity across locations* of each cell (**Fig. 1m)** was defined as average pairwise correlation of the temporal tunings in each location bin. For that calculation we included the temporal tunings in location-bins in which the activity was ≥ 0.25 relative to the *location-temporal peak.* For that analysis we included cells with significant temporal tuning and significant location tuning. To compute the percentage of location tuned neurons as a function of peak response time (**Extended Data Fig. 2i**) we took all neurons with significant temporal and location tuning and binned them according to peak response time. For each peak response time bin, we counted the percentage of neurons in that bin that had a significant location tuning out of all neurons with significant temporal and location tuning.

To compute pairwise *tuning similarity between neurons as a function of distance* (**Fig. 1n** and **Extended Data Fig. 2j-k**) we binned all neuronal pairs according to anatomical distance (lateral, axial, or Euclidian) using distance bins of 10 µm (lateral), 30 µm (axial), or 10 µm (Euclidian). For every distance bin (*d*) we computed the average Pearson correlation *r* between the tuning of neurons in that bin. For that analysis we considered temporal tuning (**Extended Data Fig. 2j**), location tuning (**Extended Data Fig. 2k**), or location-temporal tuning (**Fig. 1n**). For these three different calculations we included neurons with significant temporal, location, or location-temporal tuning, respectively. In all analysis, we included pairs of neurons with a minimal lateral distance of at least 10 µm. For analysis of tuning similarity as a function of axial distance, we included only pairs of neurons that resided withing a range of [10, 30] µm of lateral distance from each other. For the shuffle distribution we repeated the same analysis, after randomly permuting the location of each neuron in lateral and axial dimensions. The data in (**Fig. 1n** and **Extended Data Fig. 2j-k**) is presented as *mean ± s.e.m* across sessions (*n* = 53 sessions).

*Reward-outcome tuning*.

Temporal tuning and location tuning for each reward condition was computed as described in the previous section by trial-averaging the activity according to regular-reward, reward-increase, or reward-omission trials. To assess reward modulation, for each trial we computed *R* – the average activity over a time interval *t* = [-2, 5] s. We defined a neuron to be significantly modulated by reward increase by comparing the distribution of *R* for regular-reward trials *R_regular_* versus reward-increase trials *R*_*increase*_ using a two-sample Student’s t-test (at *p ≤ 0.05).* Analogously, we defined a neuron to be significantly modulated by reward omission (*reward-omission modulated neuron*) by comparing the distribution of *R* for regular-reward trials *R_regular_* versus reward-omission trials *R*_*omission*_ (at *p≤ 0.05).* Trial-averaged activity modulation by reward-increase or reward-omission was defined in the following way:

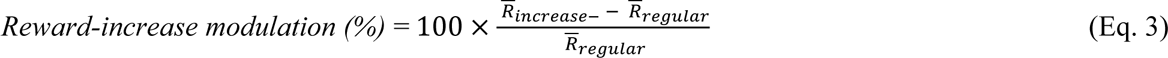

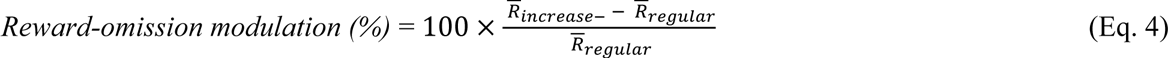

where 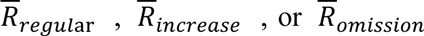 denote *R* averaged over regular-reward, reward-increase, or reward-omission trials, respectively. Note that a neuron modulated by reward-increase (or reward-omission) could be either up- or down-modulated. We refer to neurons with significant modulations as *reward-increase* or *reward-omission modulated neurons* (**Fig. 2d**).

To characterize reward-increase modulated neurons we binned them according to reward increase modulation values (**Fig. 2e**). In each bin we calculated the percentage of neurons with that level of reward-increase modulation out of all reward-modulated neurons (**Fig. 2e**, gray histogram corresponding to left y-axis). We also calculated what percentage of neurons in each bin had a significant location tuning (**Fig. 2e**, red line corresponding to the right y-axis). Analogous computation was done for reward-omission modulated neurons (**Fig. 2f**).

To compare location tuning maps computed across different reward conditions (**Fig. 2c, g-h**), we included location bins with a minimum of 3 trials per reward condition, specifically regular reward, large reward, or reward omission. These analyses included cells meeting all three of the following criteria: 1) Cells where at least 50% of their location bins had 3 trials per reward condition in each bin; 2) Cells exhibiting significant modulation in response to reward increase or reward omission; 3) Cells displaying significant location tuning. In **Fig. 2g**, we calculated, for each cell, the Pearson correlation coefficient (*r*) of location-tuning maps for regular reward versus reward-increase trials (for neurons with significant reward-increase modulation) or for regular reward versus reward-omission trials (for neurons with significant reward-omission modulation). **Fig. 2g** displays the distribution of r values for all the included cells. In **Fig. 2h** we computed the distribution of changes of location peak-width (*P*W) based on location tuning maps of different reward conditions, expressed as 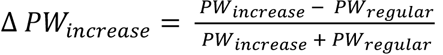 for reward-increase modulated neurons or as 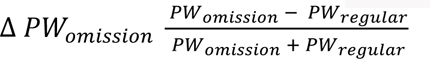 for reward omission-modulated neurons. **Fig. 2h** shows the combined distribution of Δ *P*W_*increase*_ and Δ *P*W_ommision_ values for all included neurons.

We tested if the observed modulation of neural activity by reward increase, or reward omission could be attributed to variations in lick-rate under these conditions (**Extended Data Fig. 3a**). Our approach involved several steps. First, we binned the neural activity on each trial using time bins size of 0.5 s. Then, for each trial in the data, we measured the instantaneous lick-rates at each time bin. For each time bin, we binned trials according to the instantaneous lick rate in that time bin, using lick-rate bins ranging from 0 to 12 licks/s (with a lick-rate bin size of 2 licks/s). We then trial averaged the neural activity as a function of all combinations of time along the trial and lick-rate at that time. In other words, for each neuron we created a two-dimensional tuning as a function of *time×lick-rate*. Note that the marginal distribution of this two-dimensional tuning along the time axis is the temporal tuning.

We next reconstructed the expected temporal tuning of a neuron for trials involving different reward conditions. In this simulation, we modeled the assumption that on each trial, the neural activity was solely determined by the *time×lick-rate* tuning. To achieve this, we simulated the expected neural activity for each actual behavioral trial in the dataset. This simulation involved a Poisson process with the rate parameter determined by the *time×lick-rate* tuning, corresponding to the instantaneous lick rate at each time bin within that trial. The simulated neural activity was then averaged across trials corresponding to regular-reward, reward-increase, and reward-omission conditions (**Extended Data Fig. 3c**).

We compared reward modulation (using **Eq. 3-4**) for each neuron based on real tuning and simulated tuning (**Extended Data Fig. 3d-e**). In this analysis we included neurons with significant reward-increase or reward-omission modulations. This comparison revealed that in the vast majority of neurons, reward-related modulations in the real tuning are much larger compared to those derived from lick-rate-based simulated tuning. Consequently, these findings indicate that changes in neural activity under different reward conditions cannot be trivially attributed to variations in licking behavior.

### Two-photon photostimulation protocol for causal connectivity mapping

We used an ‘all-optical’ ^5,4,6,7^ approach to measure causal connectivity. This approach involved two-photon optogenetic photostimulation of excitatory neurons and simultaneous volumetric two-photon calcium imaging of neural activity in other excitatory neurons in the field of view. Causal connectivity was assessed by detecting evoked responses in non-photostimulated neurons ^4,6^. Because the majority of neuronal Ca^2+^ influx is associated with voltage-sensitive calcium channels opened by action potentials, these responses correspond to elevated probabilities of suprathreshold responses.

Our approach involved volumetric imaging of approximately 2000 neurons while photostimulating 150 target targets in a rapidly interleaved manner, allowing rapid (∼30 mins) mapping of causal connections between ∼300,000 pairs of neurons *in vivo*. Our approach increased the number of connection pairs tested in a single experiment by more than 10-fold, compared to previous all-optical methods ^4^), as described in detail below.

The targets for two-photon photostimulation were selected with a custom MATLAB code and computer software (ScanImage) using a reference image acquired during the ‘rest only’ imaging epoch (**Fig. 3c**). The actual connectivity mapping was conducted during the ‘rest + connectivity mapping’ epoch.

We randomly picked J targets for photostimulation (typically J = ∼200 targets). Photostimulation targets were always located in the most superficial plane of the four-plane imaged volume. This is because motor cortex connectivity is directed from superficial layers to deeper layers ^44^. We photostimulated the same target while acquiring a single imaged volume and then transitioned to a new random target as we began imaging the next volume. Thus, the rate of target switching during photostimulation aligned with the rate of volumetric imaging (5.2 Hz). All targets were photostimulated in a pseudo-random order, ensuring that all J targets were photostimulated once every J trials (for a total of ∼50 photostimulation trials per target).

We stimulated specific targets with spiral photostimulation patterns ^5^. To prevent crosstalk from photostimulation into GCaMP6s imaging, we timed the photostimulation to coincide with the 15 ms gaps in acquisition between each of the four imaged planes within a volume. Spiral scans lasted 12 milliseconds, resulting in a total of 48 milliseconds of photostimulation per target (spread over the acquisition of a single imaged volume). The power of the photostimulation beam at the sample was 100–150 mW.

### Analysis of photostimulation responses

All analyses of photostimulation-induced responses were done on the deconvolved spike rates, referred to as ‘activity’ (see **Methods** section ‘Two-photon calcium imaging’). The activity of each neuron was *z*-scored. This was done to ensure that the response of each neuron to photostimulation is measured relative to its own variability and because inferring absolute levels of spike rates from calcium imaging data is challenging. We show the ΔF/F responses before deconvolution and *z*-scoring for display purpose only (**Fig. 3d** and **Extended Data Fig. 5a-c**). The change in the *z*-scored activity of the *i*^*th*^neuron upon photostimulation of the *j*^*th*^target neuron was defined as:

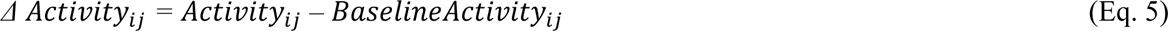

*A*_*ij*_, the activity of *i*^*th*^ neuron during photostimulation of the *j*^*th*^target, was defined as a mean activity of that neuron over a time window of 0.5 ≥ t > 0.05 s, from photostimulation onset, averaged over all trials on which the *j*^*th*^ target was photostimulated. The random sequencing of photostimulation was instrumental for isolating the response to photostimulation of individual targets by eliminating the need to wait for the activity evoked by previous random target photostimulation to revert to its baseline state. This was achieved by averaging data across trials corresponding to the photostimulation of each specific target, as described above. In addition, for each *ij* pair, we excluded trials in which nearby targets (positioned within 50 µm laterally from the *i*^*th*^ neuron) were photostimulated immediately before or after ((i.e., within ± 1 trial) the trials with photostimulation of the *j*^*th*^ target. If there were less than 20 included trials per *ij* pair, we excluded this pair from the analysis (on average we were left with 36 included trials per pair). *BaselineActivity*_*ij*_ of the *i*^*th*^ neuron was defined as the average activity of that neuron over all trials that 1) did not have a photostimulation of any target within 50 µm from the *i*^*th*^ neuron, and 2) also did not have photostimulation of nearby targets within ±1 trials from the *j*^*th*^target photostimulation.

Δ Activity_ij_ is the *causal connection strength* for any pair in which *i* ≠ *j*, or as strength of the target response to direct photostimulation for *i* = *j*. The significance of Δ Activity_ij_ was determined using one-sample Student’s *t*-test (at *p*-value < 0.05), computed for the distribution of *Δ Activity*_*ij*_ on all included individuals trials when the *j*^*th*^target was photostimulated, against the null hypothesis that the distribution has a mean equal to zero.

A neuron was ‘*directly photostimulated’* if it was within 15 µm of the photostimulation center. *‘Target neurons’* are directly-photostimulated neurons that significantly increased their activity (*p* ≤ 0.01, *t-test*) in response to direct photostimulation (**Fig. 3d**, ^6^). ‘C*ontrol targets’* are neurons that were directly photostimulated but did not show a significant activity increase (*p* > 0.1, *t-test*). In a typical experiment, J targets decomposed into ∼ 150 target neurons and ∼50 control targets (**Fig. 3e)**. Control targets were within a median distance of 53.8 µm lateral distance to target neurons (**Extended Data Fig. 6b**). Therefore, control targets were intermingled with target neurons, but simply did not respond to direct photostimulation. Control targets still had neurons above or below, as well as neurites from nearby neurons, which could potentially respond to photostimulation (**Extended Data Fig. 6**).

Neurons with significant changes in activity (*Δ Activity_ij_*) residing further away from the photostimulation center (at lateral distance > 25 µm) were considered ‘causally *connected’* to the target neuron at photostimulation center (**Fig. 3d**). We used a cutoff of 25 µm lateral distance to define causal connections because neurons at a lateral distance smaller than 20 µm from the photostimulation center could still be directly photostimulated, albeit weakly. We set this threshold by comparing the photostimulation evoked response of target neurons to that of control targets (**Fig. 3g** and **Extended Data Fig. 6**) and by characterizing the photostimulation resolution in our previous study ^6^.

Causal *connection probability as a function of distance* from the target neuron (**Fig. 3f**) was defined:

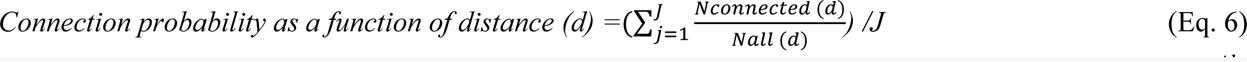

We binned all the *i*, *j* pairs according to the Euclidean distance (within the 3D imaged volume) between the *i*^*th*^ neuron and the *j*^*th*^ target, using 10 µm distance-from-target bins (*d)*. For every distance from the target bin (*d*), we computed the proportion of connected neurons *(Nconnected*) by counting the neurons that had a 1) significant and 2) positive change in activity (*Δ Activity_i,j_*), out of all neurons *(Nall)* at that bin, and averaged across all targets J. To increase the robustness of our estimation of connection probability, the photostimulation trials for every target were divided into two non-overlapping sets (odd and even). We used the odd trials to compute the significance of the connection (i.e., the significance of the change in activity, *Δ Activity*_*i*,*j*_), and the even trials to compute the sign of the connection (i.e., whether Δ *Activity*_*i*,*i*_ was positive or negative). We included only pairs with positive activity changes because we photostimulated and imaged excitatory neurons. This calculation was done separately for target neurons and control targets. For these analyses, we included sessions (*n*=24 sessions) with at least 25 target neurons and 25 control targets, photostimulated with 150 mW at sample (the power used in most experiments). The data in **Fig. 3f** is presented as *mean ± s.e.m* of connection probabilities across sessions.

*Causal connection strength as a function of distance* from the photostimulated target (**Fig. 3g**) was defined:

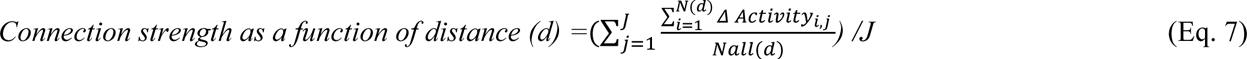

We binned all the *i*, *j* pairs according to distance (lateral, axial, or both) between the *i*^*th*^neuron and the *j*^*th*^target, using 10 µm distance-from-target bins (*d)* for lateral distance, or 30 µm for axial distance (corresponding to the distance between imaging planes). For every distance-from-target bin (*d*) we computed the average change in activity *Δ Activity*_*i*,*j*_ by summing all the *Δ Activity*_*i*,*j*_ for all neurons in that bin, divided by the number of all neurons *(Nall)* at that bin, averaged across all targets J. This calculation was done separately for directly photostimulated and control targets. Note that in contrast to connection probability calculation, which focused on pairs with significant positive change in activity (**Eq. 6**), here we included the response of all neurons, regardless of the sign or significance of *Δ Activity*. To characterize the axial profile of connection strength at different lateral distances from the photostimulation center (**Fig. 3h**), we used the same calculation as in (**Eq. 7**) but restricted the analysis to neuronal pairs residing within specific ranges of lateral distance from the photostimulation center (at lateral distances of [25, 100 µm] for Fig. 3h, left; or [100, 250 µm] for Fig3.h, right). For these analyses, we included sessions (*n*=24 sessions) with at least 25 target neurons and 25 control targets, photostimulated with 150 mW at sample (the power used in most experiments). The data in **Fig. 3g-h** is presented as *mean ± s.e.m* of connection probabilities across sessions.

To compare photostimulation evoked response on single trials (**Extended Data Fig. 5**), we categorized trials according to the strength of the evoked response (Δ Activity) of target neurons, into trials with ‘strong’ or ‘weak’ response of the target (Methods). Strong and weak trials corresponded to the 20% top and bottom percentiles, respectively, of the response (Δ Activity) of the target neuron across photostimulation trials. We show the response on strong and weak trials as ΔF/F for display purposes only (**Extended Data Fig. 5a-c**). For this calculation, responses were computed over a time window of 2 ≥ t > 0.05 s following photostimulation. We also computed the Pearson correlation (*r*) between Δ Activity of target neurons and causally connected neurons on a trial-by-trial basis, using all trials. Data in **Extended Data Fig. 5d** is shown for all neuronal pairs with significant causal connections (at *p* < 0.01; *n* = 33,830 connection pairs).

### Relating causal connectivity to functional properties

#### Causal connectivity versus noise correlations

We analyzed the relationship between causal connectivity (analogous to effective connectivity) and noise correlations (sometimes referred to as functional connectivity ^77^). We computed the pairwise noise correlation between each target neuron and the rest of the neurons with lateral pairwise distances > 25 µm. Noise correlations were defined as Pearson correlation *r* between the activity of the *j*^*th*^target neuron and the *i*^*th*^imaged neuron, with activity binned in time (1.5 s bins). Connection strength for these pairs was computed as *Δ Activity_*ij*_* (**Eq. 5**). Importantly, noise correlations were measured independently from connection strength measurements. Specifically, noise correlations were measured during either the ‘rest only’ or ‘behavior’ epochs, whereas connectivity was measured during the ‘rest + connectivity’ epoch. To relate noise correlations to connection strength, we binned all pairs according to their causal connectivity, and for each bin, we took the average noise correlation of all pairs in that bin (**Extended Data Fig. 7b-c** top, red). We included bins that had at least 50 neuronal pairs per session, from at least 5 sessions in each bin.

#### Causal connectivity versus tuning correlations

We analyzed the relationship between causal connectivity and correlations in location tuning, sometimes called signal correlations – in an analogous way to noise correlation analysis (see previous paragraph). Correlations in location tuning were defined as Pearson correlation *r* between the location tuning curves of the *j*^*th*^target neuron and the *i*^*th*^imaged neuron (**Extended Data Fig. 7a** top, red). For this analysis, we included all pairs that satisfied the following criteria: 1) 25 µm < pairwise lateral distance ≤ 100 µm (to include only pairs of cells within distance range for which there was a significant connection probability – see **Fig. 3f**); and 2) both the *i*^*th*^ and the *j*^*th*^ neurons had a significant location tuning (Methods; location tuning map stability *r* (odd, even) ≥ 0 and location-tuning modulation ≥ 25%). We included bins that had at least 25 neuronal pairs per session, from at least 5 sessions in each bin.

#### Distance-dependence of correlations and connection strength

Both causal connection strength (**Fig. 3g**) and noise correlations decrease as a function of anatomical distance between neurons ^27^. We examined if the association between connection strength and noise correlations goes beyond what can be explained by mutual distance dependence. All included pairs (as detailed in the preceding section) were binned based on their lateral-axial distance, using 10 µm lateral bins and 30 µm axial bins (the distance between planes). The noise correlation values among cell pairs in the same lateral-axial bins were randomly shuffled. In this way, the average distance-dependent profile of noise correlation remained the same in the shuffled data. However, within each distance bin in the shuffled data, there was no longer a relationship between noise correlation and connectivity strength for individual pairs – beyond what can be explained by mutual distance dependence.

The shuffled data enabled us to estimate: 1) the extent to which the relationship between connectivity strength and noise correlation is explicable by distance dependency alone, and 2) to unveil the residual relationship that remains beyond what is explained by distance dependency. To determine the *residual noise correlation, r* (**Extended Data Fig. 7b-c**, bottom), for each connectivity strength bin, we calculated the average noise correlation in that bin after subtracting the corresponding shuffled average noise correlation values (**Extended Data Fig. 7b-c**, top). These analyses were conducted separately for each session, and presented as *mean ± s.e.m* across sessions. We also did the opposite analysis by binning all pairs according to noise correlation values and computing the residual causal connectivity strength (**Extended Data Fig. 7d-e**).

Similar mutual dependency on anatomical distance exists between pairwise connection strength (**Fig. 3g**) and correlations in location tuning (**Fig. 1n**). We applied an analogous approach to uncover the relationship between connection strength and correlations in location tuning (**Fig. 4a**, **Extended Data Fig. 7a**). We repeated these analyses separately for target neurons and control targets. The *residual tuning correlation* measurement is a very conservative estimate of the relationship between tuning and connectivity strength. This is because the mutual distance-dependency between tuning correlations and connection strength (**Fig. 1n**; **Fig. 3g**) is expected under the assumptions of 1) distance-dependent synaptic connectivity ^49^, and 2) that like-to-like connectivity contributes to tuning formation ^66,2,53^.

### Network analysis of causal connectivity patterns

To perform network analysis of causal connectivity between neurons, for each target neuron, we computed its out-degree *d*_*out*_ as the number of neurons it was causally connected to within the imaged volume. For each target neuron, we also computed its in degree *d*_*in*_ as the number of neurons it receives causal connections from. *d*_*out*_ was potentially limited by the number of imaged neurons (∼2000). In contrast, *d*_*in*_ was limited by the number of target neurons (∼150 target neurons) – because we could know if a particular target receives a connection only if we directly photostimulated one of the other target neurons, which happened to be presynaptic to this particular neuron. Thus, our experimental approach was more suitable for studying outgoing (*d*_*out*_) rather than incoming (*d*_*in*_) connections, and therefore, most of our analyses were focused on out-degree connections.

For *d*_*out*_ calculations, we included only 1) significant excitatory connections – i.e., with positive change in activity; and 2) only those that were within 25 µm < lateral distance ≤ 100 µm range from the target neuron (to include only cells within distance range for which there was a significant connection probability – see **Fig. 3f**). Data in **Fig. 4c, f-h** is based on all causal connections pairs that fit the above inclusion criteria (*n* = 65,390 connection pairs; *n* = 44 sessions). When estimating *d*_*out*_ we corrected for variability in cell density around each target neuron. Specifically, we defined *relative cell density* as total number of neurons within 25 µm < lateral distance ≤ 100 µm range from the target neuron, divided by the average of this number across all targets. For example, if cell density around a particular target neuron is less than average, its *relative cell density* will be 1 <). We then divided *d*_*out*_ by the relative cell density. In all analyses we focused only on excitatory connections because we photostimulated and imaged excitatory neurons and therefore considered any inhibitory responses to be polysynaptic (whereas excitatory responses could be monosynaptic or polysynaptic). Directed causal connectivity graphs (for a representative session; **Fig. 4b**) were drawn as a top-view projection of the imaged field of view (collapsed across all imaged planes), with nodes corresponding to the location of imaged neurons. Node size indicated the number of outgoing causal connections for each target neuron (*d*_*out*_) and edges indicated connection direction and strength (**Fig. 4b**, colored arrows).

To compute the out-degree distribution (**Fig. 4c**, red), we binned the out-degree values (*d*_*out*_) for all target neurons using equally spaced bins (from 0 to maximal *d*_*out*_, using all target neurons across sessions (*n* = 44 sessions), and took the proportion of neurons in each bin out of all target neurons. We compared the out-degree distribution of a motor cortex network to that of a random network model (Erdős–Rényi), with an average degree 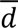 (defined as the average *d*_*out*_ across all target neurons). The degree distribution of such a random network (**Fig. 4c**, gray) follows a Poisson distribution with a rate parameter equal to 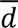 (at the limit when 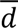 ≪ number of neurons ^78^).

For a ‘fair’ comparison between in- and out-degree connectivity, we limited our analysis to a subnetwork *S* consisting of only target neurons (**Fig. 4d-e**). For each neuron in such a subnetwork, we could fully map its causal connectivity with other neurons in the subnetwork, which we used to calculate its in-degree 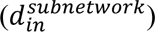 and out-degree 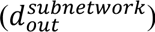. For this analysis we included all pairs with lateral distance > 25 µm. For all neurons in this subnetwork, we computed the Pearson correlations coefficients *r* between 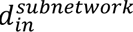 and 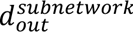 in each session. **Fig. 4d** shows the distribution of *r* across sessions (*n* = 52 sessions). We then extended this analysis to progressively more densely connected subnetworks of neurons. Specifically, we limited the subnetwork of target neurons only to those having their *d*_*out*_ above a specific minimal out-degree (*m*). For each value of *m* (ranging from 0 to 20) we computed the mean Pearson correlation *r* of the correlation between 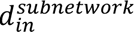 and 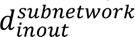 across all sessions (**Fig. 4e)**. For each value of *m*, we included only sessions that had at least 25 target neurons with *d*_*out*_ > *m* (52-18 sessions, for different values of *m*). The distribution in **Fig. 4d** corresponds to *m* = 0. Data in **Fig. 4e** is presented as *mean ± s.e.m* across sessions.

To examine the relationship between the out-degree distribution and the functional properties of neurons (**Fig. 4f-h**), we binned the out-degree (*d*_*out*_) distribution of target neurons. In **Fig. 4f**, we assessed *local noise correlation* as a function of out-degree. To achieve this, we calculated the average Pearson correlation *r* between the target neuron’s activity and the activity of each neuron within a local neighborhood. Local neighborhood was defined as all neurons within a lateral distance range of [25, 100] µm from the target neuron. To calculate noise correlations, neural activity during the ‘Rest only’ epoch was binned in time using 1.5 s time bins. For each out-degree bin, we averaged the local noise correlation values of all target neurons residing in that bin. As a control, we shuffled the local noise correlation values across all of these neurons 100 times. In **Fig. 4g**, we assessed *location tuning* as a function of out-degree. As a metric for location tuning of each neuron, we calculated the location tuning modulation (**Fig. 4g**, left) or location tuning stability (**Fig. 4g**, right; see Method section, ‘Analysis of task-related neuronal tuning’). We then averaged the values of either of these metrics across all target neurons residing in the same out-degree bin. As a control, we shuffled the location metrics of all target neurons 100 times.

Analogously, in **Fig. 4h**, we assessed *reward-outcome tuning* as a function of out-degree. To achieve this, we calculated the absolute values of reward-increase modulation and reward-omission modulation of each target neuron (**Eq. 3-4**) and took their average as an overall metric for reward-outcome modulation. We then averaged these reward modulation values across all target neurons residing in the same out-degree bin. As a control, we shuffled the reward modulation values (computed as described above) across all of these neurons 100 times.

## Data availability

Behavioral and neurophysiological data was stored and analyzed in custom pipelines in the DataJoint framework ^79^ and will be available upon request.

## Code availability

Custom MATLAB code that was used for data analysis is available on Github: https://github.com/arsenyf/2p_code_arseny

## Acknowledgments

We thank Jon Arnold and Jeff Talbot for mechanical engineering, Dmitri Tsyboulski for help with optical engineering, Sarah Lindo for help with surgical procedures, Lucas Kinsey for help with behavioral experiments, and Nuo Li for comments on the manuscript.

## Author Contributions

AF and KS designed the experiments. AF collected the experimental data with help from KD and MR. AF analyzed the data with inputs from KD, RD, and KS. AF and KS wrote the paper, with input from all the authors.

---

**Extended Data Fig. 1.**
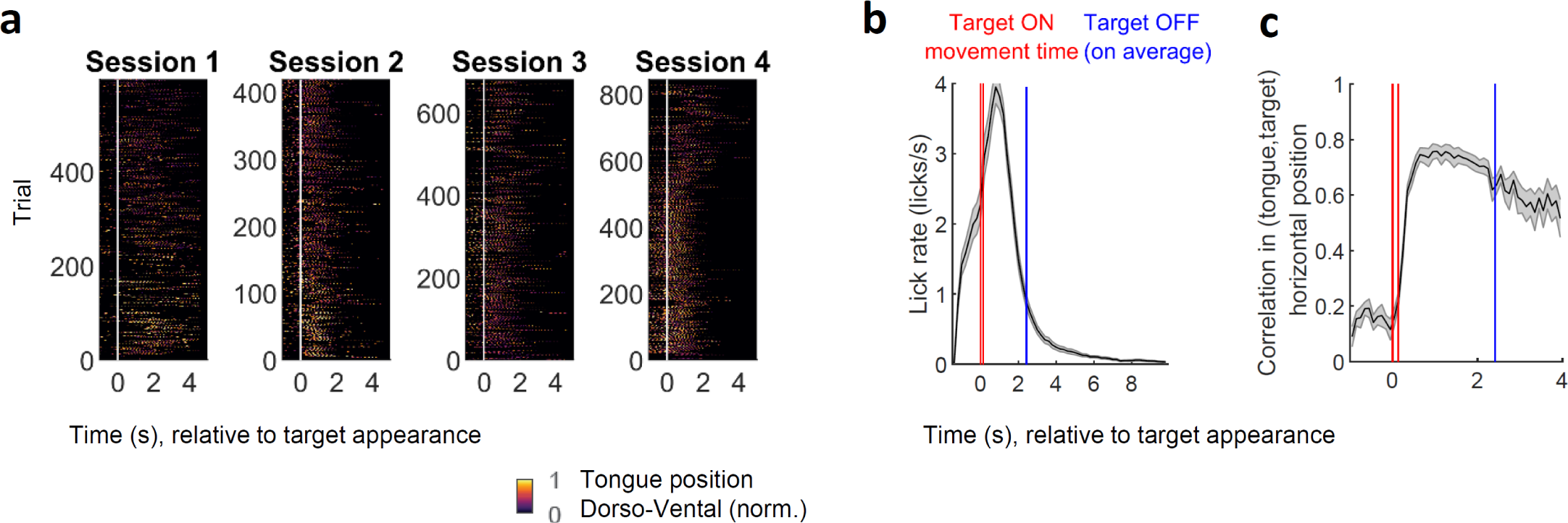
Multidirectional tongue-reaching behavior. **a,** Videography of tongue trajectories in 4 behavioral sessions of one mouse in the multidirectional tongue reaching behavior. Vertical line (white) indicates the time of target appearance. In each trial (horizontal lines) the tongue location is color-coded according to its dorsal-ventral coordinate (normalized to the maximal dorsal-ventral tongue location detected in each session). Mice produced hundreds of trials already from the first behavioral session. **b**, Lick rate as a function of time from the target appearance. Red, the interval marking the target movement towards a reachable location (‘Target ON’); Blue, average target retraction time (‘Target OFF’). Target is retracted 1 s after reward delivery or maximal time-out time (Methods), therefore its retraction time (blue) is varied on each trial. Most licks occur within less than 2 s from the target appearance. **c**, Correlation between tongue horizontal location and target horizontal location during the trial. Data in b-c is displayed as mean ± s.e.m. across sessions.

**Extended Data Fig. 2.**
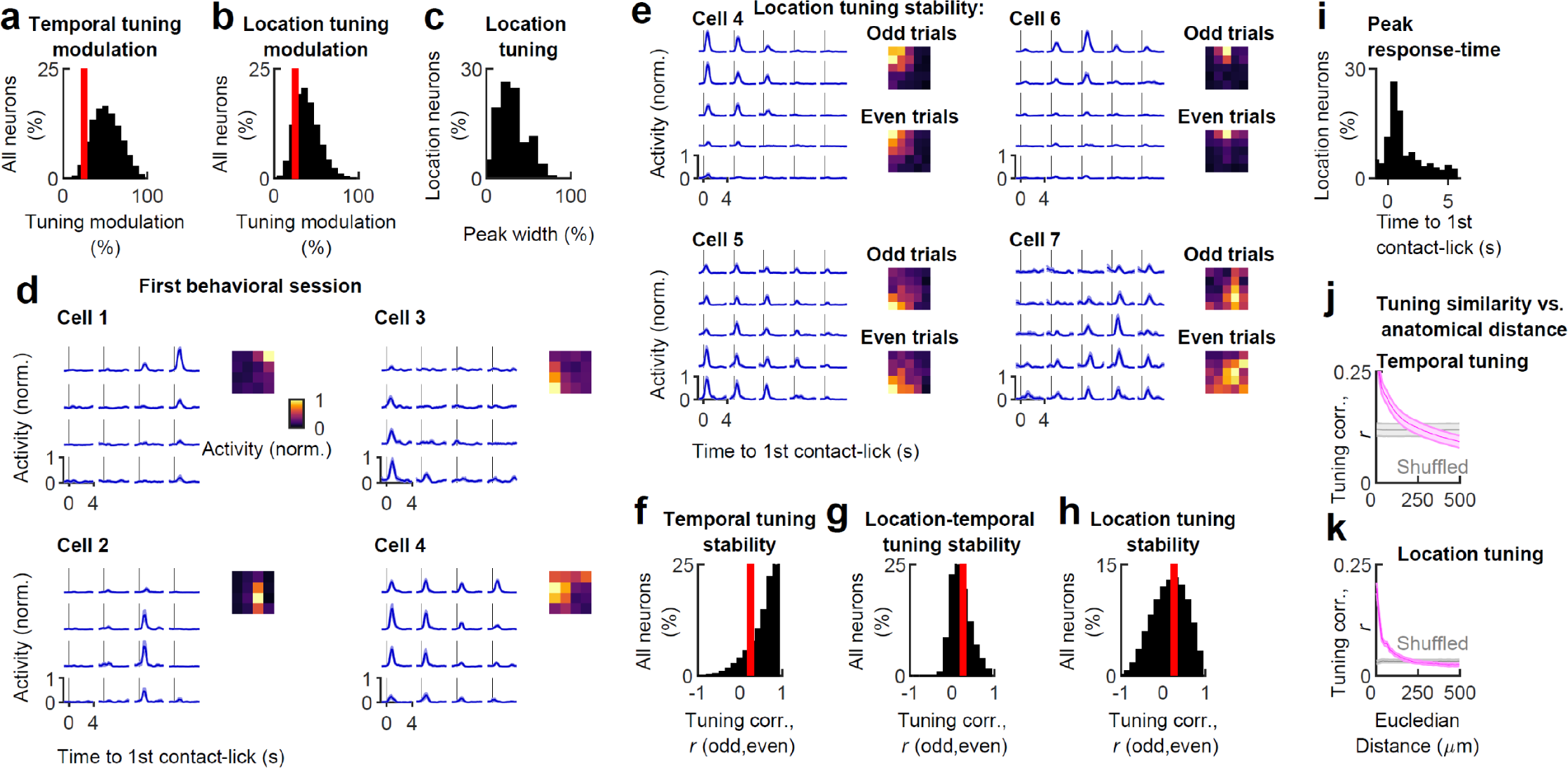
Location-temporal tuning properties. **a-b,** Distribution of temporal tuning modulation (a) and location tuning modulation (b) of all cells. Vertical lines (red; in a-b, f-h) indicate the cutoff used to categorize cells as neurons with significant tuning (Methods). **c**, Distribution of peak width of location tuning maps of location-tuned neurons. **d**, Traces showing trial-averaged responses in each target location (left) and location tuning maps (right) of 4 example cells imaged on the first day of behavior. A 4×4 location grid was used in this experiment. **e**, Location-temporal tuning of four example cells showing tuning stability. Left, traces showing trial-averaged responses in each target location. Right, location tuning map computed using two non-overlapping sets of trials (odd and even trials). A 5×5 location grid was used in this experiment. **f-h**, Distribution of stability of temporal tuning (f) and location-temporal tuning (g), and location tuning (h) of all cells computed as Pearson correlation (*r*) between tuning calculated based on odd versus even trials. **i**, Preferred peak-response time of neurons with locational tuning. **j-k,** Tuning similarity between pairs of neurons as a function of their anatomical distance for temporal tuning (j) and location tuning (k); displayed as mean ± s.e.m. across sessions.

**Extended Data Fig. 3.**
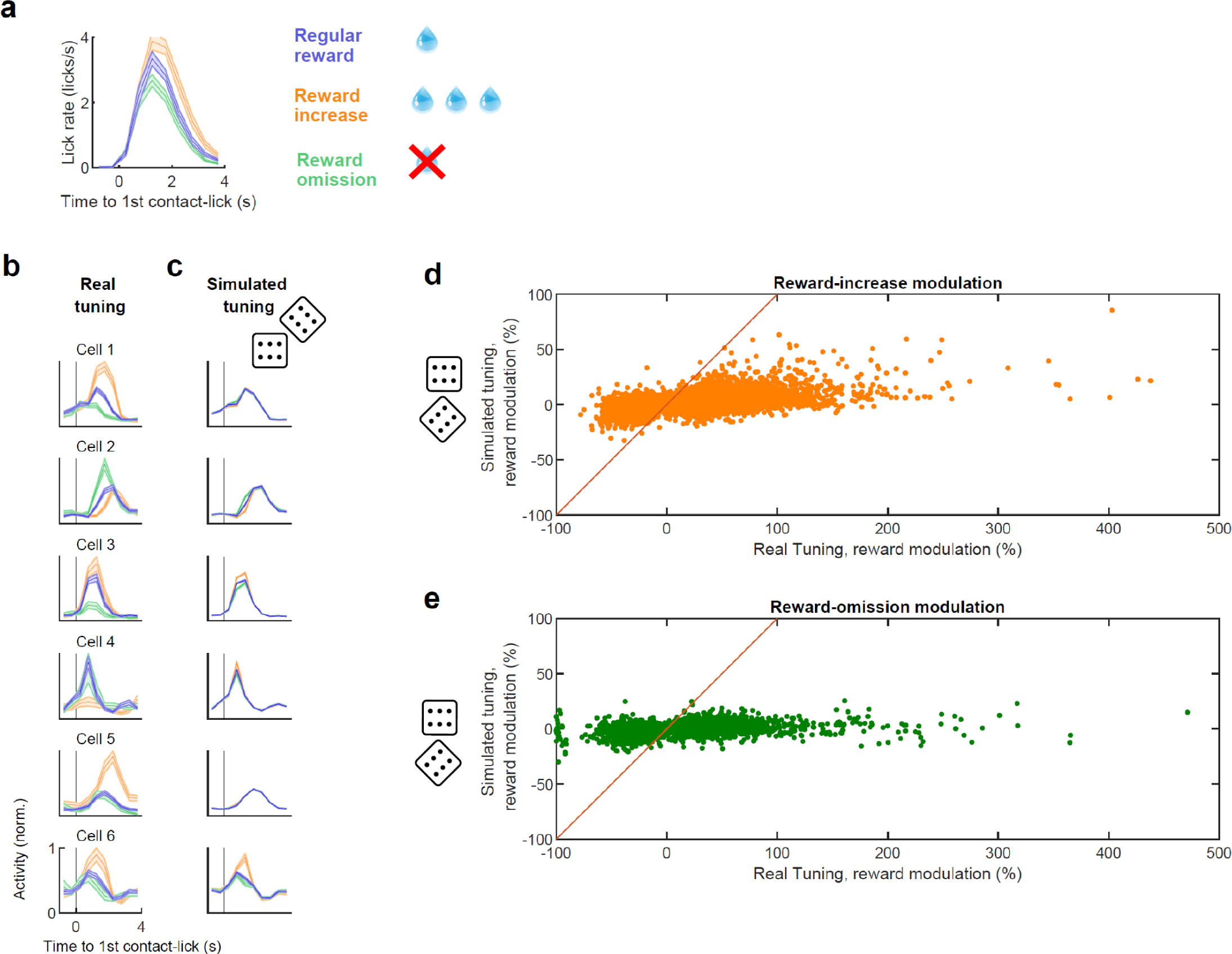
Reward-outcome modulation is not explained by variations in lick-rate. **a,** Distribution of lick-rates at different times during the trial, for reward-increase, reward-omission, or regular trials. **b**, Temporal tuning of six examples cells with reward-outcome modulation, averaged across trials with different reward conditions. **c**, Simulated temporal tuning of the same cells, generated solely based on their tuning to lick-rate at different times of the behavioral trial (Methods). Note the absence of reward modulation in simulated tuning curves (c), which suggests that reward modulation in the real data (b) is not explained by difference in lick-rates for different reward conditions for most cells (except Cell 6). **d,** Reward-increase modulation of all neurons with significant reward-increase modulation, computed based on simulated tuning versus real tuning. Each dot represents a neuron. **e,** Same as in (d) for neurons with significant reward-omission modulation. Data in a,d,e is displayed as mean ± s.e.m. across sessions. Data in b-c is displayed as mean ± s.e.m. across trials.

**Extended Data Fig. 4.**
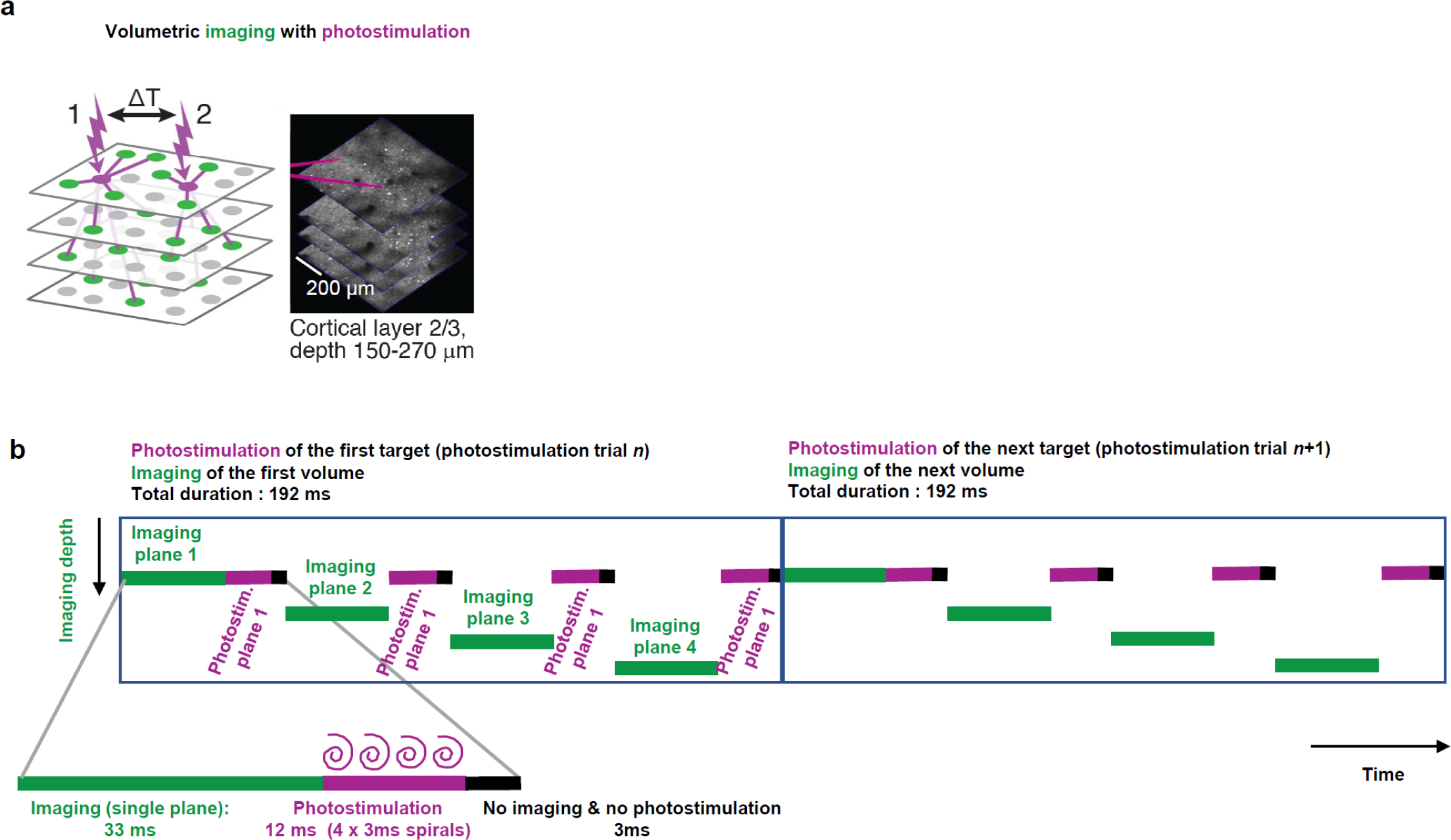
Two-photon photostimulation and volumetric imaging protocol. **a**, Left, Schematics of two-photon photostimulation of targeted neurons (magenta) and measurement of evoked responses in non-stimulated neurons (green) using volumetric calcium imaging. Right, example imaged volume of layer 2/3, motor cortex, with two photostimulation targets indicated (magenta). Both photostimulated target neurons and imaged neurons were excitatory cells. **b**, Schematics of photostimulation timing during volumetric imaging shown for two consecutive photostimulation trials. Target cells were photostimulated in the superficial plane (‘plane 1’). Imaging each plane lasted 33 ms, followed by a 15 ms gap without imaging. Four spiral photostimulations (each spiral lasting for 3 ms, see inset) occurred in the gap between each imaged plane (Methods).

**Extended Data Fig. 5.**
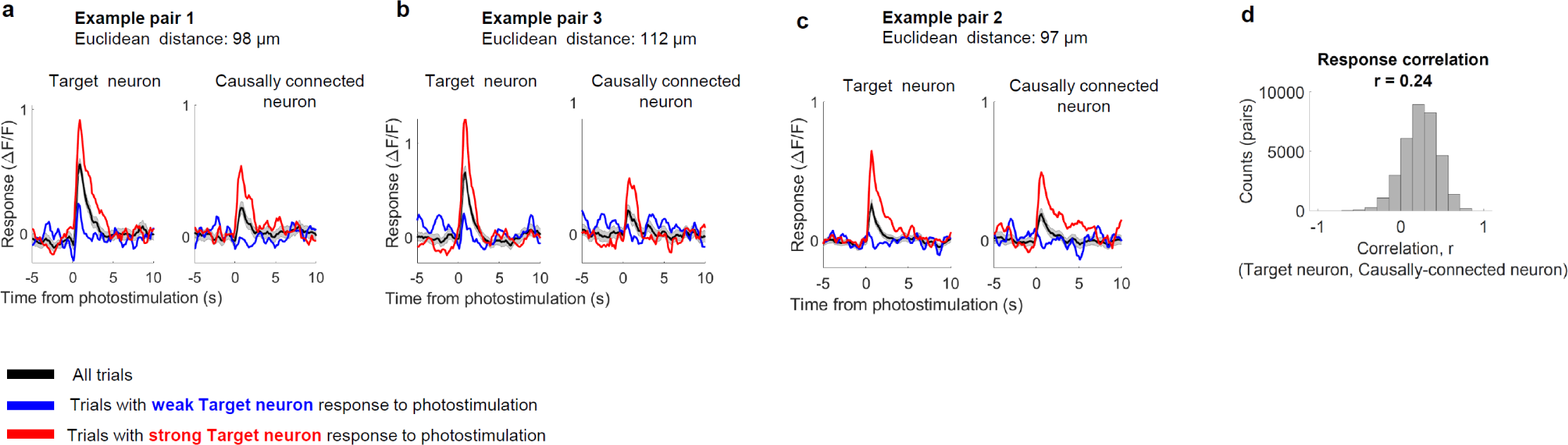
Photostimulation evoked responses on single trials. **a-c**, Comparison of the amplitude of photostimulation evoked response of the target neuron and causally connected neurons, shown for 3 example neuronal pairs. Trials were categorized according to the strength of the evoked response (Δ Activity) of the target neurons, into trials with ‘strong’ or ‘weak’ response of the target (Methods). Responses of both target and causally connected neurons are shown by trial averaging the activity using strong (red), weak (blue), or all (black) trials. **d)** Distribution of Pearson correlation (*r*) between Δ Activity of target neurons and causally connected neurons on a trial-by-trial basis, shown for all neuronal pairs with significant causal connections (*n* = 33,830 connection pairs; Methods).

**Extended Data Fig. 6.**
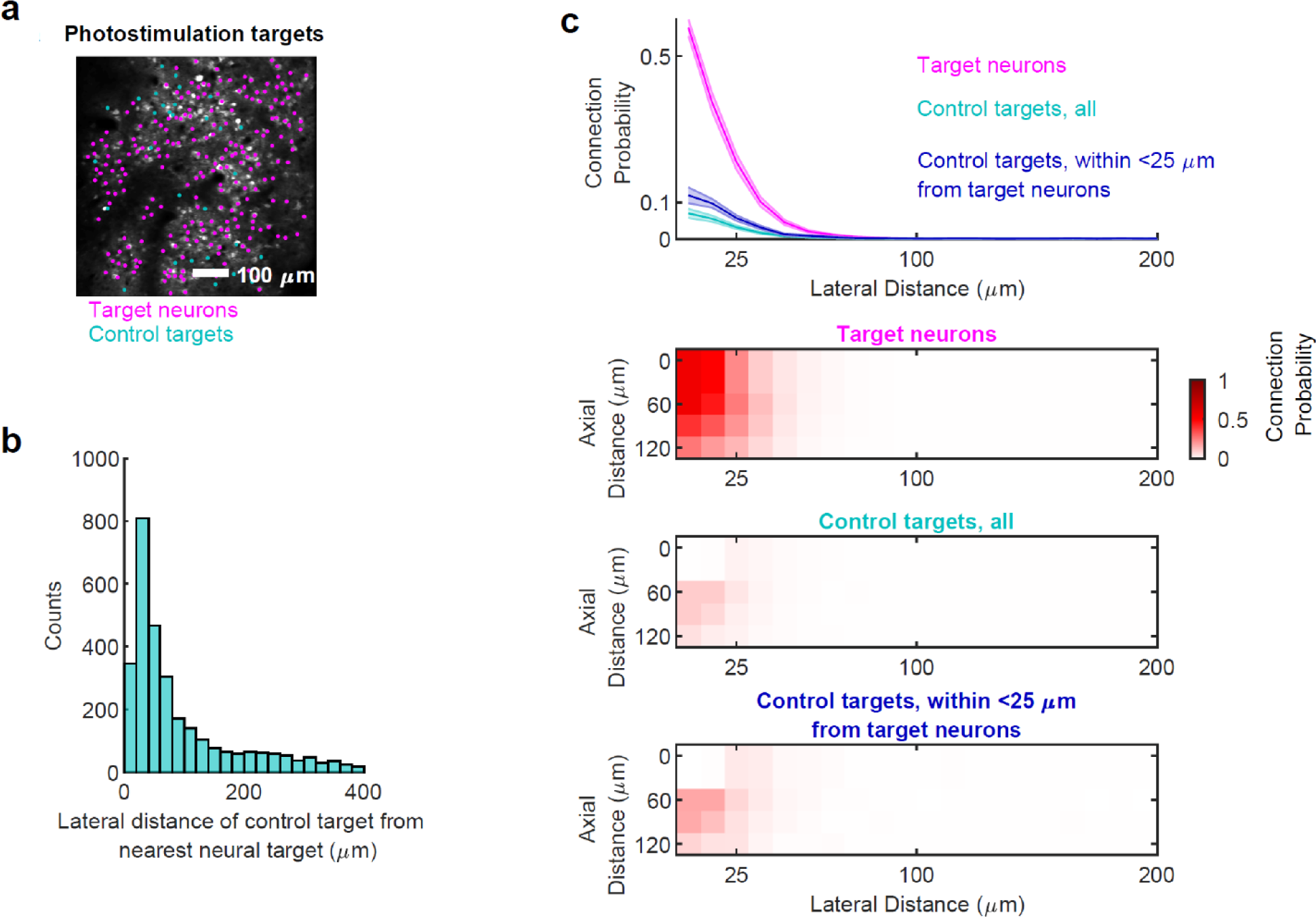
Spatial localization and causal connectivity of control and target neurons. **a,** Example field of view showing the top imaged plane with responding target neurons (magenta) and non-responding neurons referred to as ‘control targets’ (cyan). **b**, Distribution of lateral distances between each control target and the closest neuronal target for all control targets. **c,** Causal ‘connection probability’ as a function of anatomical distance (lateral and axial) between all pairs of targets and causally connected neurons, shown for target neurons (magenta), all control targets (cyan). We also analyzed connection probability only for control targets that were intermingled with target neurons (within less than 25 µm laterally from target neurons, blue). Top, display the marginal distribution of the three bottom panels as a function of lateral distance; displayed as mean ± s.e.m. across sessions.

**Extended Data Figure 7.**
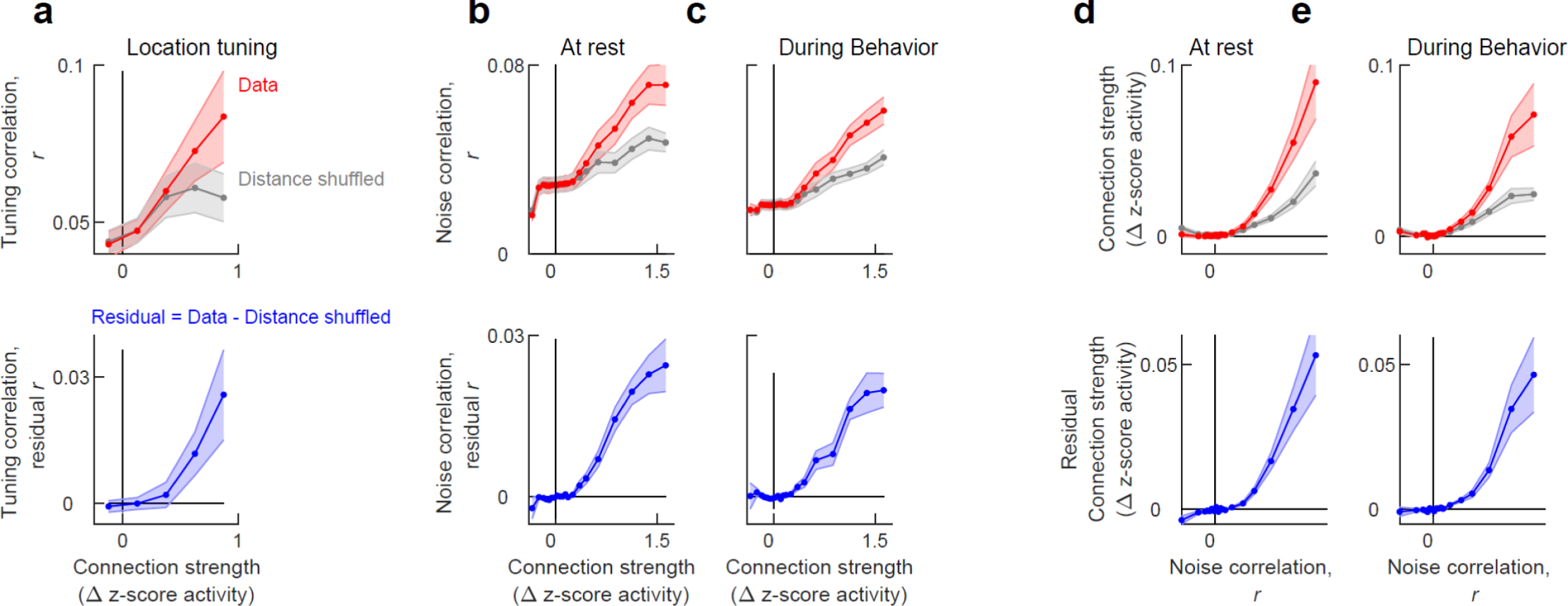
Relationship between causal connectivity and tuning or noise correlations. **a**, Comparison of causal connectivity strength with location tuning correlation measured during multidirectional reaching behavior (Methods). Top, Location-tuning correlations versus causal connectivity strength. Real data (‘Data’, red); distance-preserved shuffled control distribution (‘Distance shuffled control’, gray) represents shuffling of the correlation for all neurons residing in the same distance from the target neuron (Methods). Bottom, Residual tuning correlations versus causal connectivity strength. Residual tuning correlation (‘Residual, blue) is obtained by subtracting the ‘Distance shuffled control’ from ‘Data’ (top; Methods), and represents the residual relationship that remains between connection strength and correlation in location tuning, beyond what is explained by mutual distance dependency. **b-c**, Comparison of causal connectivity strength measured during rest, with residual ‘noise correlations’ (Methods) measured for the same neurons during rest (b) or multidirectional reaching behavior (c). Top, Noise correlations versus causal connectivity strength. Noise correlations are measured during rest (b) or multidirectional reaching behavior (c), for the same neurons. Causal connectivity is measured in an independent rest session epoch. Real data (‘Data’, red); distance-preserved shuffled control distribution (‘Distance shuffled control’, gray) represents shuffling of the correlation for all neurons residing in the same distance from the target neuron (Methods). Bottom, Residual noise correlations versus causal connectivity strength. Residual noise correlation (‘Residual, blue) is obtained by subtracting the ‘Distance shuffled control’ from ‘Data’ (top; Methods), and represents the residual relationship that remains between connection strength and noise correlations, beyond what is explained by mutual distance dependency. Causal connectivity could accurately predict noise correlations (sometimes referred as ‘functional connectivity’) across states. **d-e** Same in b-c, but for the opposite relationship, i.e., causal connectivity strength versus noise correlations. This indicates to what extent noise correlation can predict, on average, the strength of causal connectivity in the motor cortex. Data in all panels is displayed as mean ± s.e.m. across sessions.

## References

1. Hubel, D. H. & Wiesel, T. N. Receptive fields, binocular interaction and functional architecture in the cat’s visual cortex. The Journal of Physiology 160, 106–154 (1962).

2. Cossell, L. et al. Functional organization of excitatory synaptic strength in primary visual cortex. Nature 518, 399–403 (2015).

3. Yu, J., Gutnisky, D. A., Hires, S. A. & Svoboda, K. Layer 4 fast-spiking interneurons filter thalamocortical signals during active somatosensation. Nature Neuroscience 19, 1647–1657 (2016).

4. Chettih, S. N. & Harvey, C. D. Single-neuron perturbations reveal feature-specific competition in V1. Nature 567, 334 (2019).

5. Rickgauer, J. P., Deisseroth, K. & Tank, D. W. Simultaneous cellular-resolution optical perturbation and imaging of place cell firing fields. Nat Neurosci 17, 1816–1824 (2014).

6. Daie, K., Svoboda, K. & Druckmann, S. Targeted photostimulation uncovers circuit motifs supporting short-term memory. Nat Neurosci 24, 259–265 (2021).

7. Randi, F., Sharma, A. K., Dvali, S. & Leifer, A. M. Neural signal propagation atlas of C. elegans. Preprint at 10.48550/arXiv.2208.04790 (2023).

8. Bruce, C. J. & Goldberg, M. E. Primate frontal eye fields. I. Single neurons discharging before saccades. Journal of Neurophysiology 53, 603–635 (1985).

9. Churchland, M. M. et al. Neural population dynamics during reaching. Nature 487, 51–56 (2012).

10. Georgopoulos, A. P., Kalaska, J. F., Caminiti, R. & Massey, J. T. On the relations between the direction of two-dimensional arm movements and cell discharge in primate motor cortex. J. Neurosci. 2, 1527–1537 (1982).

11. Guo, Z. V. et al. Flow of Cortical Activity Underlying a Tactile Decision in Mice. Neuron 81, 179–194 (2014).

12. Peters, A. J., Chen, S. X. & Komiyama, T. Emergence of reproducible spatiotemporal activity during motor learning. Nature 510, 263–267 (2014).

13. Inagaki, H. K., Fontolan, L., Romani, S. & Svoboda, K. Discrete attractor dynamics underlies persistent activity in the frontal cortex. Nature 566, 212 (2019).

14. Finkelstein, A. et al. Attractor dynamics gate cortical information flow during decision-making. Nature Neuroscience 1–8 (2021) doi:10.1038/s41593-021-00840-6.

15. Xu, D. et al. Cortical processing of flexible and context-dependent sensorimotor sequences. Nature 603, 464–469 (2022).

16. Stepanyants, A. et al. Local Potential Connectivity in Cat Primary Visual Cortex. Cerebral Cortex 18, 13–28 (2008).

17. Schneider-Mizell, C. M. et al. Cell-type-specific inhibitory circuitry from a connectomic census of mouse visual cortex. 2023.01.23.525290 Preprint at 10.1101/2023.01.23.525290 (2023).

18. Hooks, B. M. et al. Organization of Cortical and Thalamic Input to Pyramidal Neurons in Mouse Motor Cortex. J. Neurosci. 33, 748–760 (2013).

19. Petreanu, L., Mao, T., Sternson, S. M. & Svoboda, K. The subcellular organization of neocortical excitatory connections. Nature 457, 1142–1145 (2009).

20. DeNardo, L. A., Berns, D. S., DeLoach, K. & Luo, L. Connectivity of mouse somatosensory and prefrontal cortex examined with trans-synaptic tracing. Nat Neurosci 18, 1687–1697 (2015).

21. Wu, L.-G. & Saggau, P. Presynaptic inhibition of elicited neurotransmitter release. Trends in Neurosciences 20, 204–212 (1997).

22. Marder, E. Neuromodulation of Neuronal Circuits: Back to the Future. Neuron 76, 1–11 (2012).

23. Javadzadeh, M. & Hofer, S. B. Dynamic causal communication channels between neocortical areas. Neuron 110, 2470–2483.e7 (2022).

24. Rowland, J. M. et al. Propagation of activity through the cortical hierarchy and perception are determined by neural variability. Nat Neurosci 26, 1584–1594 (2023).

25. Scott, S. H. Inconvenient Truths about neural processing in primary motor cortex. The Journal of Physiology 586, 1217–1224 (2008).

26. Tanji, J. & Evarts, E. V. Anticipatory activity of motor cortex neurons in relation to direction of an intended movement. Journal of Neurophysiology 39, 1062–1068 (1976).

27. Komiyama, T. et al. Learning-related fine-scale specificity imaged in motor cortex circuits of behaving mice. Nature 464, 1182–1186 (2010).

28. Laubach, M., Wessberg, J. & Nicolelis, M. A. L. Cortical ensemble activity increasingly predicts behaviour outcomes during learning of a motor task. Nature 405, 567–571 (2000).

29. Huber, D. et al. Multiple dynamic representations in the motor cortex during sensorimotor learning. Nature 484, 473–478 (2012).

30. Esmaeili, V. et al. Rapid suppression and sustained activation of distinct cortical regions for a delayed sensory-triggered motor response. Neuron 0, (2021).

31. Li, N., Chen, T.-W., Guo, Z. V., Gerfen, C. R. & Svoboda, K. A motor cortex circuit for motor planning and movement. Nature 519, 51–56 (2015).

32. Bollu, T. et al. Cortex-dependent corrections as the tongue reaches for and misses targets. Nature 1–6 (2021) doi:10.1038/s41586-021-03561-9.

33. Inagaki, H. K. et al. Neural Algorithms and Circuits for Motor Planning. Annual Review of Neuroscience 45, 249–271 (2022).

34. Chen, T.-W. et al. Ultrasensitive fluorescent proteins for imaging neuronal activity. Nature 499, 295–300 (2013).

35. Friedrich, J., Zhou, P. & Paninski, L. Fast online deconvolution of calcium imaging data. PLOS Computational Biology 13, e1005423 (2017).

36. Pachitariu, M. et al. Suite2p: beyond 10,000 neurons with standard two-photon microscopy. 061507 Preprint at 10.1101/061507 (2017).

37. Schultz, W., Dayan, P. & Montague, P. R. A Neural Substrate of Prediction and Reward. Science 275, 1593–1599 (1997).

38. Platt, M. L. & Glimcher, P. W. Neural correlates of decision variables in parietal cortex. Nature 400, 233–238 (1999).

39. Stuphorn, V., Taylor, T. L. & Schall, J. D. Performance monitoring by the supplementary eye field. Nature 408, 857–860 (2000).

40. Sugrue, L. P., Corrado, G. S. & Newsome, W. T. Matching Behavior and the Representation of Value in the Parietal Cortex. Science 304, 1782–1787 (2004).

41. Hirokawa, J., Vaughan, A., Masset, P., Ott, T. & Kepecs, A. Frontal cortex neuron types categorically encode single decision variables. Nature (2019) doi:10.1038/s41586-019-1816-9.

42. Pereira-Obilinovic, U., Hou, H., Svoboda, K. & Wang, X.-J. Brain mechanism of foraging: reward-dependent synaptic plasticity or neural integration of values? 2022.09.25.509030 Preprint at 10.1101/2022.09.25.509030 (2022).

43. Klapoetke, N. C. et al. Independent optical excitation of distinct neural populations. Nature Methods 11, 338–346 (2014).

44. Hooks, B. M. et al. Laminar Analysis of Excitatory Local Circuits in Vibrissal Motor and Sensory Cortical Areas. PLOS Biology 9, e1000572 (2011).

45. Mateo, C. et al. In Vivo Optogenetic Stimulation of Neocortical Excitatory Neurons Drives Brain-State-Dependent Inhibition. Current Biology 21, 1593–1602 (2011).

46. Levy, R. B. & Reyes, A. D. Spatial Profile of Excitatory and Inhibitory Synaptic Connectivity in Mouse Primary Auditory Cortex. J. Neurosci. 32, 5609–5619 (2012).

47. Sadeh, S. & Clopath, C. Theory of neuronal perturbome in cortical networks. PNAS 117, 26966–26976 (2020).

48. Dalgleish, H. W. et al. How many neurons are sufficient for perception of cortical activity? eLife 9, e58889 (2020).

49. Perin, R., Berger, T. K. & Markram, H. A synaptic organizing principle for cortical neuronal groups. PNAS 108, 5419–5424 (2011).

50. O’Connor, D. H., Peron, S. P., Huber, D. & Svoboda, K. Neural Activity in Barrel Cortex Underlying Vibrissa-Based Object Localization in Mice. Neuron 67, 1048–1061 (2010).

51. Yu, Y.-C., Bultje, R. S., Wang, X. & Shi, S.-H. Specific synapses develop preferentially among sister excitatory neurons in the neocortex. Nature 458, 501–504 (2009).

52. Das, A. & Fiete, I. R. Systematic errors in connectivity inferred from activity in strongly recurrent networks. Nat Neurosci 23, 1286–1296 (2020).

53. Khona, M. & Fiete, I. R. Attractor and integrator networks in the brain. Nat Rev Neurosci 23, 744–766 (2022).

54. Gal, E. et al. Rich cell-type-specific network topology in neocortical microcircuitry. Nature Neuroscience 20, 1004–1013 (2017).

55. Lemon, R. N. Descending Pathways in Motor Control. Annual Review of Neuroscience 31, 195–218 (2008).

56. Bari, B. A. et al. Stable Representations of Decision Variables for Flexible Behavior. Neuron 103, 922–933.e7 (2019).

57. Hattori, R., Danskin, B., Babic, Z., Mlynaryk, N. & Komiyama, T. Area-Specificity and Plasticity of History-Dependent Value Coding During Learning. Cell 177, 1858–1872.e15 (2019).

58. Lee, R. S., Engelhard, B., Witten, I. B. & Daw, N. D. A vector reward prediction error model explains dopaminergic heterogeneity. 2022.02.28.482379 Preprint at 10.1101/2022.02.28.482379 (2022).

59. Ko, H. et al. Functional specificity of local synaptic connections in neocortical networks. Nature 473, 87–91 (2011).

60. Lee, W.-C. A. et al. Anatomy and function of an excitatory network in the visual cortex. Nature 532, 370–374 (2016).

61. Bock, D. D. et al. Network anatomy and in vivo physiology of visual cortical neurons. Nature 471, 177–182 (2011).

62. Kuan, A. T., et al. Synaptic wiring motifs in posterior parietal cortex support decision-making. http://biorxiv.org/lookup/doi/10.1101/2022.04.13.488176 (2022) doi:10.1101/2022.04.13.488176.

63. Ginzburg, I. & Sompolinsky, H. Theory of correlations in stochastic neural networks. Phys. Rev. E 50, 3171–3191 (1994).

64. Oldenburg, I. A. et al. The logic of recurrent circuits in the primary visual cortex. 2022.09.20.508739 Preprint at 10.1101/2022.09.20.508739 (2022).

65. Amsalem, O., Inagaki, H., Yu, J., Svoboda, K. & Darshan, R. Sub-threshold neuronal activity and the dynamical regime of cerebral cortex. 2022.07.14.500004 Preprint at 10.1101/2022.07.14.500004 (2022).

66. Ben-Yishai, R., Bar-Or, R. L. & Sompolinsky, H. Theory of orientation tuning in visual cortex. Proceedings of the National Academy of Sciences 92, 3844–3848 (1995).

67. McNaughton, B. L., Battaglia, F. P., Jensen, O., Moser, E. I. & Moser, M.-B. Path integration and the neural basis of the ‘cognitive map’. Nature Reviews Neuroscience 7, 663–678 (2006).

68. Guo, Z. V. et al. Maintenance of persistent activity in a frontal thalamocortical loop. Nature 545, 181–186 (2017).

69. Chen, S. et al. Brain-wide neural activity underlying memory-guided movement. 2023.03.01.530520 Preprint at 10.1101/2023.03.01.530520 (2023).

70. Song, S., Sjöström, P. J., Reigl, M., Nelson, S. & Chklovskii, D. B. Highly Nonrandom Features of Synaptic Connectivity in Local Cortical Circuits. PLoS Biology 3, e68 (2005).

71. Towlson, E. K., Vértes, P. E., Ahnert, S. E., Schafer, W. R. & Bullmore, E. T. The Rich Club of the C. elegans Neuronal Connectome. J. Neurosci. 33, 6380–6387 (2013).

72. Uzel, K., Kato, S. & Zimmer, M. A set of hub neurons and non-local connectivity features support global brain dynamics in C. elegans. Current Biology 32, 3443–3459.e8 (2022).

73. Bonifazi, P. et al. GABAergic Hub Neurons Orchestrate Synchrony in Developing Hippocampal Networks. Science 326, 1419–1424 (2009).

74. Bollmann, Y. et al. Prominent in vivo influence of single interneurons in the developing barrel cortex. Nat Neurosci 26, 1555–1565 (2023).

75. Daie, K., et al. ALM Window Surgery. (2021).

76. Mathis, A. et al. DeepLabCut: markerless pose estimation of user-defined body parts with deep learning. Nat Neurosci 21, 1281–1289 (2018).

77. Friston, K. J. Functional and effective connectivity in neuroimaging: A synthesis. Human Brain Mapping 2, 56–78 (1994).

78. Fundamentals of Brain Network Analysis. (Academic Press, 2016). doi:10.1016/B978-0-12-407908-3.09996-9.

79. Yatsenko, D. et al. DataJoint: managing big scientific data using MATLAB or Python. bioRxiv 031658 (2015) doi:10.1101/031658.

